# Developmental conditions shape lifetime reproductive strategies in a wild mammal

**DOI:** 10.64898/2025.12.05.692614

**Authors:** Lauren Petrullo, David Delaney, Stan Boutin, Jeffrey E. Lane, Andrew G. McAdam, Ben Dantzer

## Abstract

Coping with developmental hardship is a defining feature of resilience across the tree of life. Compensatory strategies that preserve fitness after a poor start are hypothesized to be favored by natural selection, but empirical evidence is rare. Using over three decades of data on >1,000 reproductive attempts across the lifetimes of 405 wild female red squirrels, we show that developmental adversity prompts adaptive reorganization of reproductive investment across the lifespan. Hardships reduced lifetime reproductive opportunities, but triggered opposing strategies. Extrinsic mortality risk predicted frontloading of reproduction at debut, while poor maternal investment predicted terminal reproductive effort at a cost to future generations. Both strategies preserved lifetime fitness, revealing how context-dependent reproductive plasticity over the life course can rescue the fitness consequences of developmental hardship.

## Main Text

For nearly a century, evolutionary biologists have probed the lifetime costs of harsh developmental conditions (1–4). Organisms that experience early-life adversity are not expected to accrue the same fitness as individuals born in better quality environments [i.e., the “silver spoon effect” or developmental constraints, (5–7)]. Empirical data in nonhuman mammals have largely supported this hypothesis: early-life hardship hampers later reproductive success across species (8–10). In humans, adversity is thought to similarly reduce future reproductive potential through negative effects on reproductive maturation and reproductive physiology (11, 12). Harsh conditions can become biologically embedded in developing individuals through life history trade-offs that preserve short-term survival but exacerbate cellular aging and inflammation (13), constraining reproductive success and reducing lifetime fitness (14).

Reproductive strategies that mitigate the fitness costs of early-life hardship have been proposed [e.g., (15–18)], and are hypothesized to be favored by natural selection (19–21). When poor developmental conditions cue similarly poor conditions in adulthood, anticipatory adjustments that match future extrinsic mortality risk or later-life somatic condition may minimize reproductive costs (22). For example, accelerating reproductive maturity or redistributing reproductive effort sooner in life could allow individuals with poor survival prospects to begin accruing reproductive success more rapidly, offsetting the lifetime fitness costs of developmental constraints (23). Alternatively, a later-life bump in reproductive performance akin to senescence-induced terminal investment could preserve fitness if it allows developing individuals to prioritize, and perhaps even optimize, short-term condition over early reproduction (24).

Such adaptive responses may ultimately hinge on the type of adversity and its associated constraints (20). Population-level adversities that cue extrinsic mortality risk across entire cohorts of individuals [e.g., predation or competition, (25, 26)] may induce strategies that prioritize immediate accrual of fitness gains to preserve geometric mean fitness (27). By contrast, adversities related to maternal investment that reduce condition (e.g., sibling competition, prematurity) may prompt individual-level adjustments to reproductive timing that minimize long-term somatic damage, preserving relative fitness by maintaining competitive ability (20, 28). Although these types of adversities are not mutually exclusive [e.g., extrinsic mortality can cause somatic damage, (29)], their distinction can help disentangle whether adversity-induced constraints are consequences of harsh conditions themselves, or rather byproducts of otherwise compensatory strategies.

Yet, empirical evidence for adaptive responses to developmental hardship is sparse (28, 30, 31). A reliance on fitness proxies rather than reproductive success, a tendency to focus on single sources of adversity, and a lack of longitudinal data on reproductive effort across the entire life course have limited our ability to distinguish adaptive responses from assumed costs. A historical focus on long-lived species has further obscured potential compensatory strategies because selective pressures favoring redistribution of reproductive effort are expected to be stronger in shorter-lived species with fewer reproductive opportunities (32). And fluctuating environments, which favor the evolution of phenotypic plasticity (33, 34), are more likely than those with unpredictable random change (e.g., hurricanes, lifestyle transitions) to generate enough reproductive variation on which natural selection can act to shape adaptive responses to developmental hardship (33).

### Quantifying developmental adversity in a fluctuating environment

In this study, we use 36 years of data across the lifetimes of 405 female red squirrels over >1,000 reproductive attempts over, to test the hypothesis that developmental adversity differentially cues adaptive reorganization of reproduction across the lifetime. In the Yukon, juvenile red squirrels (Tamiasciurus hudsonicus) inhabit a highly fluctuating environment in which multiple co-occurring sources of adversity activate physiological stress responses (35) and reduce short-term survival (36). We first quantified sources of developmental adversity by identifying maternal, social, and ecological variables that reduced the probability that a juvenile survived its first winter [>200 days of age, the main survival bottleneck in this system, (37)]. We used a broad, population-wide dataset (N=3,689 juveniles across 36 years) to avoid conditioning early-adversity indices on later outcomes. From this model, we extracted estimates for significant predictors of juvenile survival, finding quantitative support for birth date, litter size, the rate of postnatal growth, food availability, population density, and predation risk as sources of early-life adversity (Fig. 1A, table S1, fig. s1).

**Figure 1.**
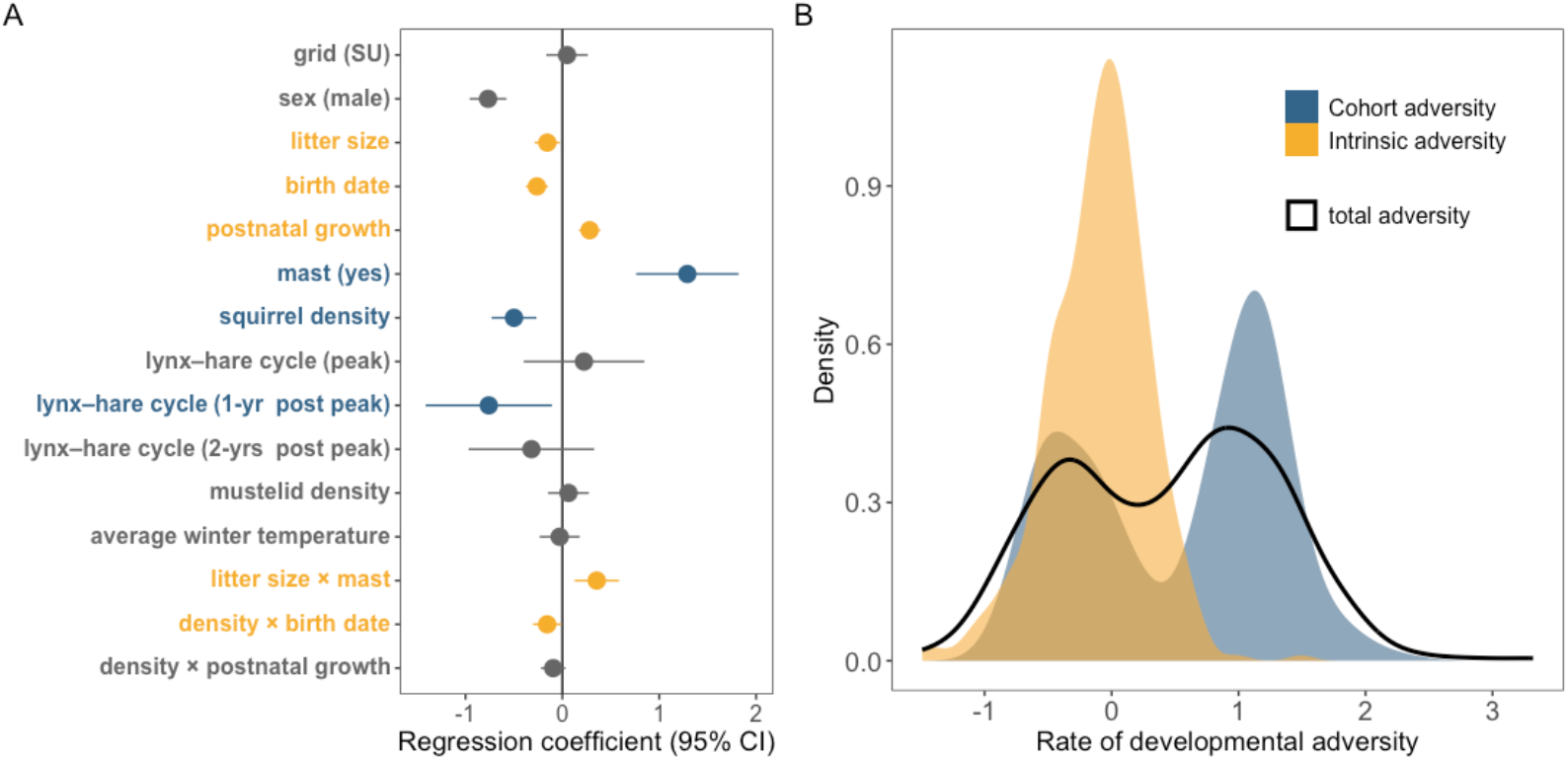
Distribution and rates of cumulative developmental adversity in wild red squirrels. **(A)** Maternal investment (orange terms) and extrinsic mortality risk (blue terms) predict variation in the probability of juvenile survival over the first winter (i.e., >200 days, table s1). We considered significant terms to be sources of early-life adversity, and extracted estimates from these terms to construct indices of developmental adversity [see Materials and Methods, (*36, 65*)]. **(B)** Cohort adversity indices reached higher magnitudes compared to intrinsic adversity indices, and were bimodal. This bimodality likely reflects the fluctuating nature of the three environmental variables that comprised this index: squirrel population density, episodic production of their primary food source (masting white spruce, *Picea glauca*), and the lynx-hare predator-prey cycle, in which the post-peak crash in snowshoe hares (*Lepus americanus*) causes Canada lynx *(Lynx canadensis*) to prey-switch to squirrels (*35*).

We used these estimates to build an undifferentiated and weighted cumulative index of total developmental adversity for females with full life course data on reproductive effort [N=405 females, see materials and methods, (36)]. We then partitioned adversities related to poor maternal investment at the individual level (i.e., intrinsic adversity), and adversities related to population-level extrinsic mortality (i.e., cohort adversity) that reduced the likelihood of juvenile survival (Fig. 1B). Cohort adversity was composed of food scarcity (i.e., a non-mast year), increased squirrel density and thus conspecific competition, and heightened predation risk. Intrinsic adversity included a slower rate of postnatal growth, being born into a large litter, and having a birth date later in the season. In general, rates of cohort adversity reached greater magnitudes than intrinsic adversity [i.e., cohort adversity predictors had larger effect sizes on average, table S1]. The two adversity indices were only weakly correlated, capturing largely distinct dimensions of early-life conditions (r=−0.07; cov=−0.022).

### Adversity reduces survival without a cost to lifetime fitness

Despite their small size, female squirrels adopt relatively “slow” life histories, investing strategically in reproduction to preserve longevity (38). Reductions to total lifespan are a commonly documented cost of early-life adversity in humans and other animals [e.g., (36, 39)], and thus represent the broadest constraint on the opportunity to accrue fitness. Across our dataset, the median adult lifespan was ∼3 years (range: 1-8, fig. s2a). Adult lifespan strongly predicted both lifetime reproductive output (total number of offspring produced) and lifetime reproductive success (LRS, total number of offspring surviving >200 days, Fig. 1A, table S2).

As developmental adversity increased, females lost nearly 2 months off their adult lifespans (β=- 0.05, P=0.007, table S3). This reduction was driven entirely by cohort adversity, which predicted both a shorter lifespan (β=-0.08, P<0.0001, table S3) and an increased hazard of death (HR=1.14, CI: 1.03–1.26, table S4). Yet, no measure of early-life adversity explained variation in lifetime fitness (table S2), or the likelihood that a female would achieve any reproductive success at all during her lifetime (i.e., LRS>0, table S5). Unlike studies documenting fitness costs of early-life adversity [e.g., (8–10)], females that experienced developmental hardship did not suffer reduced lifetime fitness in any dimension. Moreover, preserving longevity did not enhance LRS for females that experienced intrinsic adversity (β=-0.04, P=0.01; table S6), indicating that maintaining lifespan is not a universal route toward enhancing lifetime fitness. Instead, the decoupling of longevity and LRS by early-life adversity suggests alternative reproductive strategies to maintain lifetime fitness.

### Adversity compresses reproductive tenure by shifting reproductive timing

We next tested whether adversity predicted variation in a more direct measure of reproductive opportunity: the duration of reproductive tenure (i.e., time between first and last reproduction, fig. s2b). A longer reproductive tenure predicted both greater lifetime reproductive output (β=0.39, P<0.0001) and success (LRS, β=0.45, P=<0.0001, Fig. 2A), and did so more strongly than adult lifespan (Table S7). However, developmental adversity predicted compression of reproductive tenure, with females losing ∼5% with each unit increase in intrinsic adversity and ∼6% with each unit increase in cohort adversity (table S8, Fig. 2B-C).

**Figure 2.**
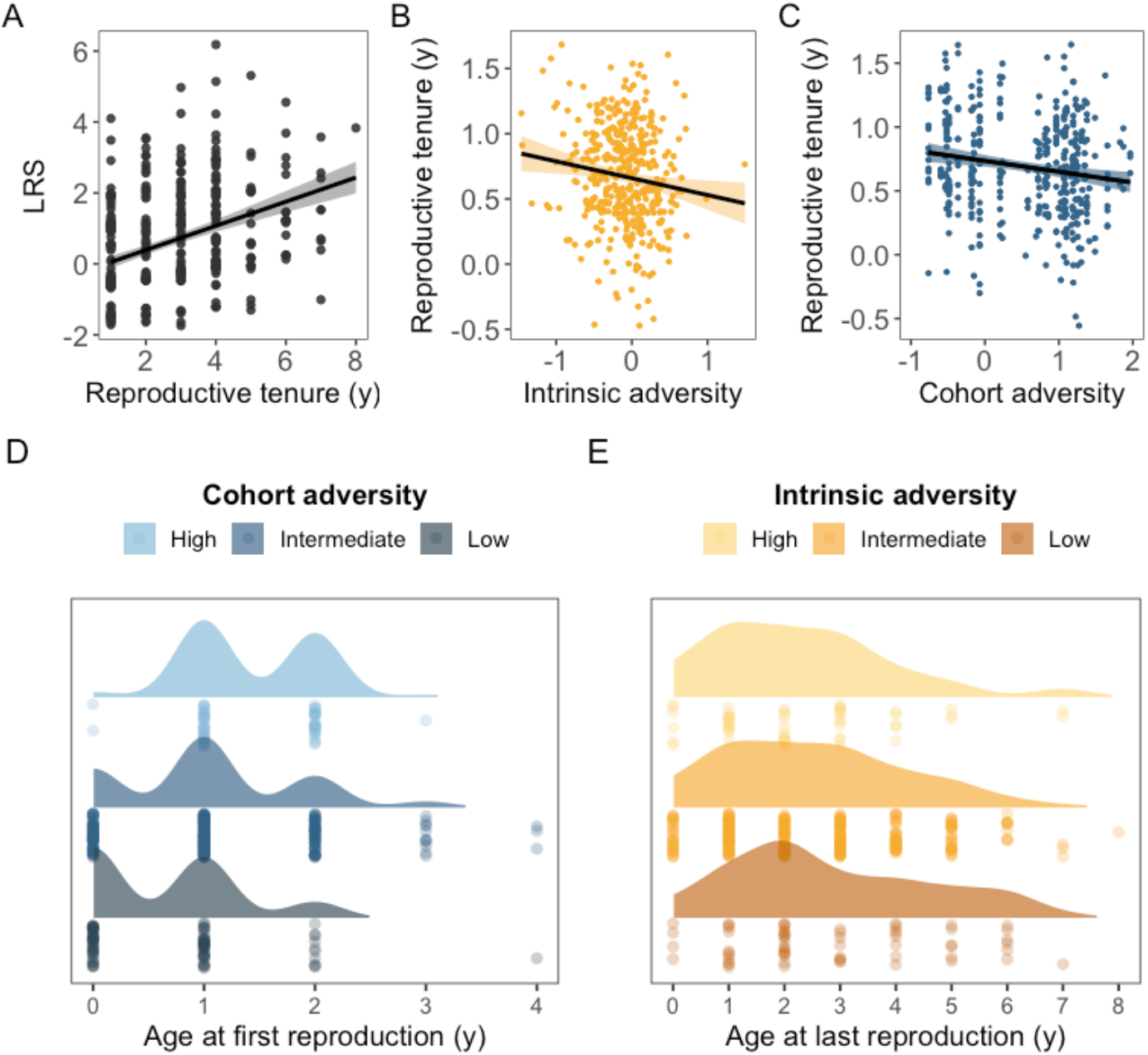
Adversity differentially abbreviates reproductive tenure. **(A)** The duration of a female’s reproductive tenure positively predicted her total lifetime reproductive success (LRS), with longer tenures resulting in a greater number of offspring surviving >200 days. However, **(B)** intrinsic adversity and **(C)** cohort adversity compressed the period of reproductive tenure (table s8). **(D)** Among females with high rates of cohort adversity, reproductive tenure was compressed by later age at first reproduction; **(E)** among females with high rates of intrinsic adversity, tenure was compressed by an earlier age at last reproduction (table s9). Scatterplots depict partial residuals from mixed-effects models controlling for total lifespan and environmental quality across the life course. Rainclouds and ridgelines depict distributions of raw data on reproductive schedules.

We hypothesized that females constrained by developmental adversity may preserve lifetime fitness by redistributing reproductive effort earlier in life (23, 28). In mammals, selection typically favors earlier reproductive maturity, especially when population sizes fluctuate and fitness benefits compound (40). Indeed, an earlier age at first reproduction predicted greater lifetime fitness across all females (table S9). However, those that experienced developmental adversity exhibited delayed, rather than accelerated, reproductive maturity. This effect was driven entirely by cohort adversity, such that age at first reproduction lagged by over two months for each standard deviation increase in cohort adversity (β=0.35, P=0.003, Fig. 2D, table S10).

Despite theoretical expectations that developmental adversity will accelerate reproductive maturity [e.g., (23)], increased predation risk and food scarcity can instead stall or entirely suspend reproductive investment as females disinvest in reproduction (41, 42). Because periods of high population density (a source of cohort adversity) trigger adaptive accelerated juvenile growth in this population (43), faster growth may generate tradeoffs by deprioritizing reproductive maturity in favor of mass gain (43, 44). Alternatively, delayed reproductive maturity could itself be an adaptive strategy to maintain competitiveness (45). If female squirrels born into years with heightened extrinsic mortality risk suffer substantial competitive deficits, they may benefit from delaying reproduction until they can recoup deficits and improve competitive ability. By affecting all individuals born in a given year, cohort adversities may homogenize among-individual variation in competitive ability, reducing selective pressure to accelerate reproduction (46).

For females that experienced intrinsic adversity, the compression of reproductive tenure was driven not by a delay in age at first reproduction but by an earlier age at last reproduction [i.e., they stopped reproducing sooner, table S11, Fig. 2E]. Because maternal investment varies among individuals within cohorts, these females may prioritize early reproductive competence to maintain competitiveness with their cohortmates as well with older individuals in the breeding population.

### Extrinsic mortality risk triggers adaptive frontloading of reproduction

When population density limits juvenile recruitment more than adult survival, selection is expected to favor rapid, risk-tolerant resource allocation decisions (47, 48). Heightened extrinsic mortality risk could thus induce a dual-phase response by both delaying reproductive maturity and increasing reproductive rate (49). We quantified female reproductive effort (i.e., number of offspring produced and rate of offspring survival) in the first breeding season as a function of developmental adversity. Controlling for the quality of the environment at reproductive debut, females with higher rates of cohort adversity produced more offspring in their first breeding season compared to females with lower rates (first-season output, β=0.13, P<0.000, table S12, Fig. 3A). Their rate of offspring survival was also higher (first-season success, β=0.29, P=0.001, table S12, Figure 3B).

**Figure 3.**
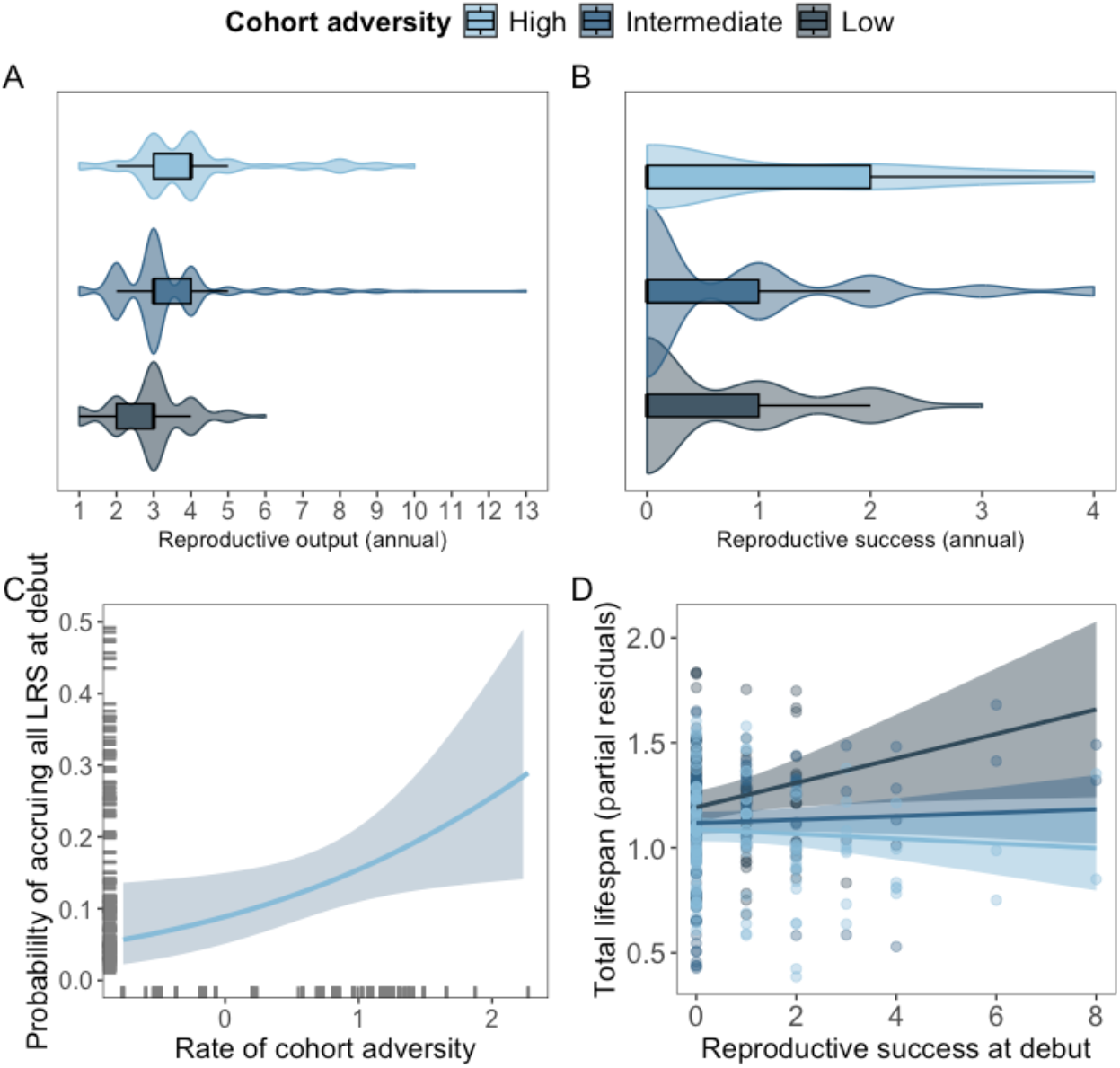
Cohort adversity predicts greater reproductive effort at the first reproductive attempt, increasing lifetime fitness. **(A)** Reproductive output and **(B)** reproductive success at debut increases with an increasing rate of cohort adversity (table s12). Adversity values displayed at three representative levels: low (−1 SD), intermediate (mean), and high (+1 SD) relative to the population distribution. **(C)** As the rate of cohort adversity increases (x-axis), females become more likely to accrue all of their lifetime fitness at reproductive debut in their first breeding season (y-axis), controlling for total lifespan (table s13). **(D)** Females with high rates of cohort adversity that frontload reproduction into the first breeding season (light blue line) suffer longevity costs (y-axis).

As cohort adversity increased, females became more likely to accrue the entirety of their LRS at reproductive debut (controlling for lifespan, β=1.51, P<0.0006, table S13, Fig. 3C). Among females that spread LRS across multiple seasons, higher rates of cohort adversity predicted a greater proportion of their total LRS accrued in the first season (β=0.62, P=0.008, table S13). By frontloading reproduction in this manner, females that experienced cohort adversity enhanced their LRS (cohort adversity × first season reproductive success; β=0.09, P=0.036, table S14, Fig. 3D). This strategy revealed a previously inconspicuous lifetime fitness penalty for females that experienced cohort adversity but did not frontload reproduction (β=-0.13, P=0.003, table S14).

Redistributing reproductive effort into the first breeding attempt could preserve geometric mean fitness for cohorts of individuals born into bad years. By reducing the multiplicative demographic costs of developmental stress across generations, desynchronized reproductive schedules may stabilize population viability, particularly in variable environments (27). We thus quantified each female’s individual generation time as her offspring-weighted mean age at reproduction (50). Despite delayed reproductive maturity, cohort adversity shortened generation times (β=-0.80, P<0.000, table S15), suggesting a faster reproductive tempo characterized by more reproduction sooner–consistent with an accelerated pace of life.

Frontloading reproduction may be especially advantageous in fluctuating environments where females can exploit favorable but rare breeding conditions. In the Yukon, red squirrels inhabit a boom-bust ecosystem in which availability of their primary food source, white spruce (Picea glauca), peaks only once every 3-7 years during mast years (51). In mast years, squirrels have the rare opportunity to dramatically boost short- and long-term fitness. Selection favors heightened responsiveness to mast cues (51, 52), yet because individuals live only ∼3 years, many die without ever experiencing a mast (∼40% in our dataset). We therefore hypothesized that females constrained by developmental adversity would show heightened sensitivity to mast cues and stronger reproductive responses to a mast at first reproduction. Cohort adversity, which includes being born in a food scarce non-mast year, predicted a higher probability of encountering a mast in the future, and a shorter time-to-mast (table s16). However, females that experienced cohort adversity were not more likely to first reproduce in a mast year (table s17), indicating that they do not delay reproductive maturity until detecting a high-payoff context. Instead, they invested heavily in reproductive output at debut regardless of mast conditions (table s18).

Principles of life history theory predict that frontloading reproduction will incur costs (53). Indeed, females that concentrated reproductive effort into the first attempt suffered steeper declines in both adult lifespan (β=-0.14, P=0.008) and reproductive tenure (β=-0.08, P=0.005) compared to those that did not (table s19). These costs were more severe if they first reproduced in a mast year (table s20), indicating that high-payoff environments intensify both reproductive responses and their associated costs. At the mechanistic level, the oxidative stress and accelerated cellular aging associated with increased reproductive rates can erode longevity, particularly for females already constrained by early-life adversity (54). However, by improving the likelihood of offspring survival, the nutritional surplus of a mast year may partially offset transgenerational effects of adversity-induced constraints from females who frontload to their offspring.

### Poor maternal investment cues terminal reproductive effort

Unlike cohort adversity, intrinsic adversity shifted reproductive investment later in life–a paced strategy more consistent with condition-driven models [e.g., internal predictive adaptive response hypothesis, (20)]. Intrinsic adversity elongated generation times (table s15), revealing a delayed reproductive strategy despite no delay in reproductive maturity. Although historically applied in the context of aging (16, 24), terminal investment–a bump in reproductive effort near death–may allow these females to prioritize recovering condition during early life while preserving lifetime fitness. Across all females, the rate of reproductive success increased linearly with proximity to death, independent of age (linear time to death; β=0.65, P<0.001; table s21). This relationship was further shaped by a negative quadratic effect, such that reproductive success peaked at intermediate proximity to death before declining (quadratic time to death; β=–0.48, P=0.001; table s21).

Females that experienced intrinsic adversity exhibited steeper linear increases in reproductive success as they approached death (intrinsic adversity × linear years-to-death; β=0.27, P=0.05, table s21), consistent with improving condition. They also exhibited later and more pronounced peaks, followed by a sharper post-peak decline (intrinsic adversity × quadratic years-to-death; β=–0.30, P=0.02; table s21; Fig. 4A). This steeper approach toward more prominent and terminal reproductive peaks suggests a form of terminal investment cued not by senescence or immunological aging, but by developmental hardship (24). By increasing reproductive output at the terminal reproductive attempt, females that experienced intrinsic adversity enhanced LRS (final season output × intrinsic adversity; β=0.02, P=0.02; table s22, Fig. 4B). Despite payoffs of this later-life reproductive investment, terminal effort was shifted toward offspring quantity rather than quality: these females were poorer at converting terminal output into surviving offspring (final season reproductive success × intrinsic adversity, table s23).

**Figure 4.**
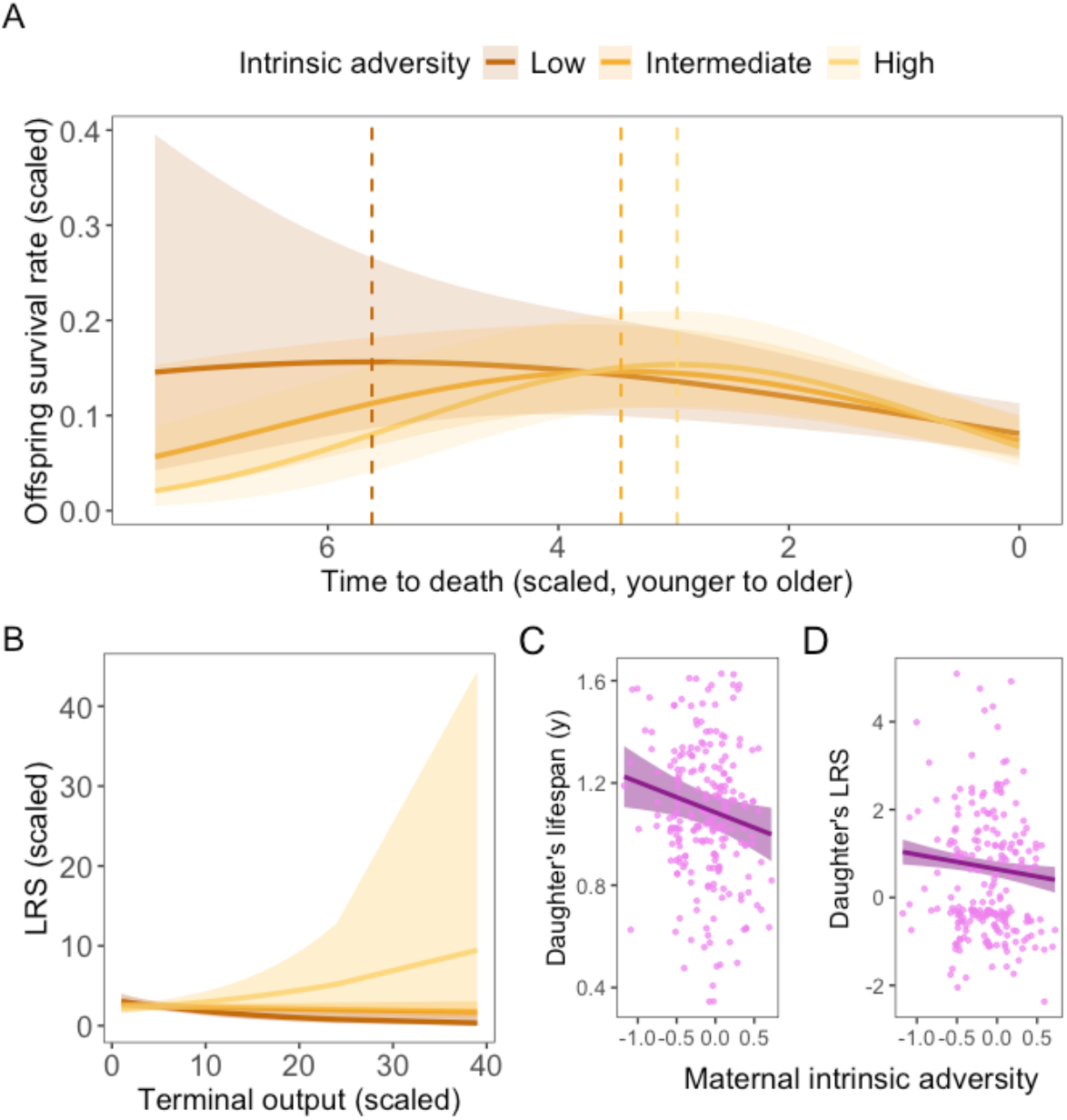
Intrinsic adversity shifts reproductive effort later in life at a cost to daughters. Females that experienced **(A)** high rates of intrinsic adversity (≥1 SD from the mean, light orange line) exhibited later reproductive peaks (dashed line indicates quadratic peak), as well as a faster ramp up to the peak and a steeper decline following the peak (table s21). **(B)** In the terminal breeding season specifically, females that experienced high rates of intrinsic adversity increased lifetime fitness by producing more offspring (table s22). Despite mothers maintaining their own lifetime fitness, daughters suffered **(C)** reduced lifespan and **(D)** poorer reproductive success, regardless of their own early-life adversity (table s24).

Poor maternal investment in the first year of life may lead to persistent physiological deficits [e.g., immune or metabolic dysfunction, (55)] that constrain later-life maternal care. Consequently, females may be able to capitalize on existing mechanistic architecture to increase reproductive output (56, 57), but remain unable to invest in offspring during the more energetically expensive period of lactation (58). We find that such constraints propagate across generations: daughters of mothers that experienced intrinsic adversity suffered poorer lifetime fitness and reduced longevity, regardless of their own experiences of adversity (N=230 mother-offspring pairs, Fig. 4C-D, table s24).

By pushing reproductive peaks later and ending their reproductive career and its associated costs sooner, females that experienced intrinsic adversity may protect future longevity while passing adversity costs to their offspring (59–62). Alternatively, poorer early-life growth, a component of intrinsic adversity, may negate a lifespan cost by promoting longevity, consistent with growth–lifespan tradeoffs (63). Ultimately, accelerated reproductive senescence may be the currency paid for a terminal strategy that protects both early-life condition and lifetime fitness in the face of poor maternal investment.

Developmental hardship can force individuals to prioritize short-term survival over other biological processes, leading to tradeoffs that constrain adaptation and evolution (64). These constraints have prompted longstanding debate over if and how organisms can compensate for the seemingly inevitable fitness costs of early-life hardship (15, 36). Here, we show that early-life adversity does not universally impose fitness costs, despite developmental constraints. Instead, females differentially shift reproductive investment over the life course. Adversities that reflect population-level extrinsic mortality risk prompted frontloading of reproductive effort at debut, which could circumvent the compounded costs of lost future reproductive opportunity. By contrast, individual-level adversities related to poor maternal investment predicted reproductive restraint and a paced strategy of terminal investment, which may protect relative fitness by improving competitive ability near death.

Collectively, our results demonstrate that, as a singular phenomenon, developmental hardship generates opposing strategies that adaptively reorganize reproductive effort across the lifespan. The costs commonly attributed to early-life adversity may therefore be, at least in part, byproducts of conditionally adaptive reproductive strategies that preserve an individual’s lifetime fitness.

## Acknowledgments

We thank Agnes MacDonald and her family for long-term access to her trapline, and the Champagne and Aishihik First Nations for allowing us to conduct our work within their traditional territory. We would especially like to thank the many field technicians that contributed to data collection, as well as Charley Krebs and the Community Ecological Monitoring Program for their continued collection of the predator-prey data used in this manuscript. This is KRSP paper #XXX.

## Funding

This work was supported by funds from the University of Arizona, the University of Michigan, the National Science Foundation (DEB-2010726 to LP, DEB-0515849 to AGM, IOS-1749627 to BD, and DEB-2338394 to AGM and BD) and the Natural Sciences and Engineering Research Council of Canada to (SB, AGM, JEL).

## Author contributions

Conceptualization: LP; Methodology: LP, DMD, AGM; Visualization: LP; Formal analysis: LP; Funding acquisition: SB, JEL, AGM, BD, LP; Project administration: SB, JEL, AGM, BD, LP; Writing – original draft: LP; Writing – review & editing: LP, DMD, SB, JEL, AGM, BD.

## Competing interests

The authors declare that they have no competing interests.

## List of supplementary Materials

Materials and Methods

Figs. S1-S2

Tables S1-S24

## Notes

### Competing Interest Statement

The authors have declared no competing interest.

