## Supplemental Materials for "Developmental conditions shape lifetime reproductive strategies in a wild mammal"

### Materials and Methods

Ethics statement

All approvals in support of this work were granted by the University of Arizona and the University of Michigan Institutional Animal Care and Use Committees (Arizona: 2024-1312, Michigan: PRO00009804, PRO00007805 and PRO00005866), and met the Canadian Council on Animal Care Guidelines and Policies. All relevant guidelines were followed. Fieldwork was permitted under Yukon Territorial Government Wildlife Research Permits and Scientists & Explorer's Permits.

Study site and population

The Kluane Red Squirrel Project (KRSP) is a long-term field study of North American red squirrels (*Tamiasciurus hudsonicus*)–highly territorial, solitary, and arboreal rodents– in the southwest Yukon, Canada. KRSP has been monitoring red squirrels in this region continuously since 1989 (*65*). Data for this study were collected over the course of 36 years (1989-2024) from wild red squirrels (total, N=405 individual lifetimes) living on one of two unmanipulated study areas ("Kloo"/KL, "Sulphur"/SU). These study areas are similar in size (~40 ha), in close proximity to one another, and experience similar environmental conditions (*35*).

Data collection

**Growth rate.** We calculated rates of postnatal growth from 0 to 25 days of age, a period of linear growth during which pups are entirely dependent on maternal milk (*66*). We temporarily removed pups from their nests for weighing within ~0 days of birth, and again at ~25 d old. Growth rates were then calculated as the gain in mass between the two nest entries (g/day). Faster growth across this postnatal period in this population consistently predicts better short-term survival [e.g., (*36*, *43*)].

**Litter size.** We counted litter sizes within 24 hours of birth. We have previously found that litter size trades off against postnatal growth, such that offspring born into larger litters are more likely to exhibit growth deficits due to sibling competition and limited maternal resources (*67*). However, in mast years, the abundance of food available during the period of juvenile independence largely ameliorates this trade-off (*67*). We therefore included an interaction between litter size and food availability (mast/non-mast year) in our preliminary model on juvenile survival.

**Birth date.** Birth dates were determined using radiotelemetry on gestating females. We used observed mating chases as approximate dates of conception, and searched for nests approximately 35 days later. In a typical year, earlier birth dates predict better juvenile survival because the competition for vacant territories is lower (i.e., there are more vacant territories earlier in the year). By contrast, later birth dates predict poorer survival because fewer vacant territories are available. When squirrel population densities are high, this effect is magnified, thus we included the interaction between birth date and squirrel density in our juvenile survival model (*67*).

**Population density.** We calculated adult squirrel population densities as the annual number of adults present on each of our study areas expressed as squirrels per hectare. Because adult dispersal is low and site fidelity is high and all individuals are uniquely tagged, we can completely enumerate the number of adult squirrels living on our study areas each year using live-trapping and behavioral observations.

**Predation risk.** We used population monitoring data on a terrestrial predator of red squirrels in this region, Canada lynx (*Lynx canadensis*), from the Kluane Boreal Forest Ecosystem Project (1987 through 1996) and the Community Ecological Monitoring Program (from 1996 onward) (*68*). We have witnessed terrestrial predation of squirrels by lynx at our field site and routinely collect data on lynx abundance, thus we chose to focus on lynx in this study. We have also previously shown that predation pressure from Canada lynx influences early-life survival (*35–37*). We used lynx snowtrack counts collected across transects during winter to calculate predator abundance index for each year (average snow track counts / 100 km transects), and camera trap data from 2020 onward. We also considered the abundance of the primary non-squirrel prey of Canada lynx: snowshoe hares (*Lepus americanus*); however, when hare densities are low, lynx exhibit prey switching to red squirrels (*68*). Hare density estimates were generated using mark-recapture. We additionally included data on the abundance of potential nest predators of red squirrels (mustelids) using transect data and snow track counts.

**Temperature data.** To assess overwinter temperatures, we obtained daily temperature records from the Haines Junction weather station (Climate ID 2100630, 60.77°N, 137.57°W), located ~35 km SE of our study area. We calculated the annual average overwinter temperature as the mean temperature across the months of October to the following March.

**Food abundance.** In this region, red squirrels primarily consume seed from masting white spruce (*Picea glauca)* trees, which produce a superabundance of food in the autumn (mid-August) episodically every 3–7 years in mast years, and little to no food in the years between (non-mast years, [39]). Across the 36 years of this study, seven (7) mast years occurred (cone distributions described in previous studies, [39,46]). In all years (mast and non-mast), we counted the number of visible cones on one side of the top third of a consistent subset of trees (between 159 and 254 trees) on each study area [66]. We then log (+1) transformed counts and calculated the mean to represent an annual index [60]. We defined years with a superabundance of cones as mast years, which occur once every 3–7 years [43].

**Annual reproductive success.** We calculated annual reproductive success as the total number of pups produced in a given breeding season that ultimately survived to at least 200 d of age (i.e., into the following breeding season). This represents a critical survival bottleneck in this population as it hinges on juvenile squirrels acquiring their own territories prior to winter, which is necessary to recruit successfully into the breeding population (*37*).

**Lifetime reproductive output and lifetime reproductive success.** We calculated lifetime reproductive output as the total number of offspring produced across the lifespan. We calculated lifetime reproductive success as the total number of offspring recruited into the breeding population (i.e., juveniles that survived >200 days over their first winter and into the following breeding season).

**Total adult lifespan.** Lifespan was determined only for females whose birth dates were known and who had died during the course of this study. We resolved lifespans to whole years (integer) to avoid conditioning longevity on the March and September dates of our population-wide censuses.

**Reproductive tenure and spread.** For all females that expressed full adult lifespans (i.e., were born and died during the course of this study), we created a full sequence of all years in which a female bred, including any years she may have been alive but skipped breeding. We quantified the duration of reproductive tenure as the time (in years) between a female's age at first reproduction and age at last reproduction (inclusive). To determine reproductive spread, we quantified reproductive output (total number of offspring produced) and reproductive success (i.e., offspring surviving >200 days) for all yearly reproductive attempts between a female's first and last breeding seasons. If a female bred, then skipped breeding the following year, but then resumed breeding, her reproductive output/success in the year she skipped was assigned zero.

**Food production and social density over the lifetime and across reproduction.** We summed the total amount of spruce cones produced across a female's total lifespan (lifetime spruce cones) as well as the total number of cones produced specifically across the specific period of her reproductive tenure (spruce cones across reproduction). For conspecific density, we summed the total number of squirrels living on a female's study area across her entire lifespan (lifetime squirrel density) as well as the total number of squirrels across the specific period of her reproductive tenure (density across reproduction). We included these variables to control for potential confounding effects of continued adversity into adulthood, as well as potential ameliorating effects of high quality future environments (*36*).

Statistical approaches

**Quantifying juvenile survival.** We defined juvenile survival as overwinter survival past 200 days of age into the following spring breeding season. This stage represents the main survival bottleneck in this system (*37*). To determine juvenile survival, we completely enumerate the number of squirrels in our study areas twice a year during study-wide censuses. In September of each year, we use behavioral observations and live-trapping to determine which juveniles born earlier that year have successfully acquired a territory. The following March, we census all squirrels again. Squirrels unaccounted for in the spring census are presumed dead with high confidence (*25*, *38*).

**Constructing early-life adversity indices.** Previous studies have suggested that cumulative indices of early-life adversity capture additive effects of developmental conditions on later-life phenotype better than single-variable models (*36*, *39*). Moreover, *weighted* indices that consider the relative effect sizes of individual sources of adversity can outperform both single-variable and cumulative count models because they account for heterogeneity in the predictive power of different types of early-life adversity (*36*). We have previously shown in our study population that a weighted early-life adversity index accounting for the relative effect sizes of different variables on juvenile survival outperforms a raw count/integer index of cumulative adversity (*36*).

To calculate a weighted early-life adversity index for each individual in our dataset, we first constructed a generalized linear mixed-effects model (family=binomial) testing the effects of covariates on the probability of overwinter survival in the first year of life (*36*). We then extracted the effect sizes for significant covariates and used these to create a total weighted early-life adversity index, as well as partitioning covariates to create specific indices for intrinsic and cohort adversity. All indices incorporated both the count and magnitude of adversities an individual experienced, as well as their continuous values. Specifically, for each significant predictor of juvenile survival (growth rate, litter size, parturition date, squirrel density, predation risk, and food availability), we extracted the regression coefficient for each from the juvenile survival model (multiplied by −1 such that positive values reflect increased adversity) and multiplied it with the value of the environment or life-history trait for each squirrel. For significant interactions, we multiplied the value of the environment experienced with each individual's life-history trait value and the interaction coefficient. Note that interactions were incorporated into intrinsic adversity indices. We then summed the strength of each axis of the environment to represent a cumulative index of early-life adversity.

**Effects of adversity on lifespan and mortality risk.** We modeled the effect of early-life adversity on total adult lifespan and mortality risk in two ways. First, we built a generalized linear model with adult lifespan as the dependent variable to determine how adversity explained variation in lifespan controlling for other covariates. Second, we build a semiparametric Cox proportional hazards model, which estimates how predictors influence mortality risk without requiring a specified baseline hazard function. The model included standardized measures of intrinsic and cohort adversity, lifetime density and lifetime food availability, whether the individual ever encountered a mast or not, and study area as fixed effects. Because all individuals had known lifespans, we assumed no right-censoring (all events = 1).

**Variation in lifetime fitness.** We modeled lifetime reproductive output using generalized linear models (GLMs) with a Poisson error distribution and log link. The first model tested whether lifetime reproductive output (total number of offspring produced) varied with developmental adversity, lifespan, and environmental covariates. The second model examined whether lifetime reproductive success (LRS; total number of offspring produced) varied as a function of the same predictors, while including the log of total lifetime reproductive output as an offset to account for differences in reproductive opportunity. Both models included measures of intrinsic and cohort adversity, lifetime density and food availability, mast experience, and study area as fixed effects.

**Variation in reproductive tenure.** We modeled reproductive tenure using generalized linear models with a Gamma error distribution and log link first with total developmental adversity as a predictor, and then with cohort and intrinsic adversity partitioned. Models tested whether developmental adversity predicted shorter reproductive tenures after accounting for total lifespan (included as a log-transformed offset). Environmental covariates during the reproductive period—squirrel density, cone availability, and mast experience—were included as fixed effects along with study area. We ran models including total adversity and separate indices of intrinsic and cohort adversity to assess their independent contributions to reproductive tenure.

We then compared the relative importance of reproductive tenure versus total adult lifespan in predicting both lifetime reproductive output and lifetime reproductive success (LRS). Using Poisson generalized linear models with a log link, we modeled lifetime reproductive output and LRS as functions of either reproductive lifespan or total lifespan, controlling for lifetime food availability, population density, mast experience, and study area. We used AICc values to evaluate model support and determine which measure (total adult lifespan vs. reproductive tenure) better explained variation in lifetime fitness.

**Variation in reproductive timing.** We modeled how variation in reproductive timing influenced reproductive tenure and lifetime fitness. We built generalized linear models with a Gamma error distribution to test how age at first (AFR) and last reproduction (ALR) predicted reproductive tenure after accounting for lifespan differences and other covariates. To assess downstream fitness effects, we fit Poisson mixed models with random intercepts for the year of first reproduction to test how AFR and ALR predicted lifetime reproductive output and lifetime reproductive success. Finally, we modeled AFR and ALR themselves as outcomes using mixed models to evaluate whether developmental adversity (total, intrinsic, or cohort) predicted delayed or truncated reproductive schedules, controlling for lifespan, environmental conditions, and study area.

**Reproductive frontloading.** To test whether early-life conditions predicted greater reproductive effort at the first reproductive attempt (i.e., frontloading), we quantified reproductive output (number of offspring produced) and reproductive success (number of offspring surviving > 200 days) in the first breeding season. We then fit two generalized linear mixed-effects models (Poisson), including intrinsic and cohort adversity, age, study area, and first-season environmental covariates (mast/non-mast year, squirrel density, and cone availability) as fixed effects. We additionally included a random effect of year of first reproduction to account for any outstanding inter-annual variation in environmental conditions not captured by our fixed effects.

Next, we evaluated whether cohort adversity increased the likelihood that females concentrated their total lifetime reproductive success (LRS) in their first breeding season. Using generalized linear mixed-effects models with binomial errors, we modeled both the binary probability of accruing all lifetime reproduction at debut and the proportion of total LRS produced in the first season. Both models included first-season year as a random effect and controlled for adult lifespan, lifetime reproductive environment, mast experience, and study area.

To test if and how frontloading as a reproductive strategy influenced lifetime fitness, we modeled total lifetime reproductive success (LRS) as a function of first-season reproductive success, cohort adversity, and their interaction using a Poisson GLM. We included reproductive lifespan as an offset to account for variation in reproductive opportunity among females. Finally, to evaluate potential survival costs of early investment, we modeled adult lifespan (Gamma with log link) as a function of cohort adversity, first-season reproductive success, and their interaction.

**Reproductive senescence and terminal investment.** We evaluated whether females exhibited terminal investment—a bump in reproductive effort near death—and whether this pattern was modulated by early-life adversity. To characterize within-individual changes in reproductive performance near death, we modeled the probability of offspring recruitment (offspring survival > 200 days) for 1,075 reproductive attempts using a generalized linear mixed-effects model with a binomial error distribution and logit link. The model included linear and quadratic terms for years from death to capture curvilinear trajectories in reproductive success, as well as interactions with intrinsic and cohort adversity to test whether early-life adversity altered the shape or magnitude of late-life reproductive investment. We also controlled for female age. Random intercepts for individual ID and study area × year accounted for repeated measures and spatial–temporal variation. Because age and proximity to death were weakly correlated (r = -0.29), we further tested whether the observed pattern of reproductive investment with increasing proximity to death could be explained by age-related reproductive trajectories alone. We compared age-structured and time-to-death-structured models to evaluate whether reproductive investment increased specifically as death approached (terminal investment) or simply followed the expected age-related peak in reproductive performance. Adversity only emerged as a significant predictor in the proximity model, and this model explained more of the variance than the age-structured model (delta AIC = ~21).

To assess how adversity influenced the fitness returns of later-life reproduction, we next modeled lifetime reproductive success (LRS) as a function of reproduction in the final breeding season. We built a generalized linear model (Poisson family, log link, offset = lifespan) testing whether females with high intrinsic or cohort adversity converted reproductive output into fewer surviving offspring at the end of life, indicating reduced efficiency of late-life reproduction. Fixed effects included age at last reproduction, lifetime reproductive output, mast experience, and environmental covariates describing food availability and squirrel density across reproductive years. Finally, to determine whether increased reproductive output in the final season compensated for this reduced efficiency, we modeled LRS as a function of late-life reproductive output, adversity measures, and their interactions. This model tested the hypothesis that females experiencing high intrinsic adversity maintained lifetime fitness through increased offspring quantity rather than quality—consistent with terminal investment.

**Individual generation time**

To quantify reproductive tempo across the life course in an integrative way, we calculated individual generation times for each female as the offspring-weighted mean age at reproduction (*50*). Specifically, for each female, the mean age at a given reproductive attempt was weighted by the total reproductive output (i.e., offspring number) at that attempt, such that reproductive attempts in which more offspring were produced exhibited proportionally greater influence. We then built a generalized linear model (Gamma family, log link) to test the hypothesis that early-life adversity affects the tempo of reproductive effort across the life course. The model included generation time as a dependent variable and fixed predictors of intrinsic adversity, cohort adversity, age at first reproduction, reproductive tenure, lifetime spruce cones, lifetime squirrel density, experienced mast (yes/no), and study area.

**Intergenerational effects**

To test whether early-life adversity experienced by one generation influences fitness outcomes in the next, we constructed a two-generation dataset linking each female in our dataset to her mother, such that we had full lifespan data for 230 mother-daughter pairs. We then tested for intergenerational effects of both types of adversity. We first modeled daughter's adult lifespan with a Gamma error distribution and log link as a function of her own early-life adversity (cohort and intrinsic indices) as well as her mother's early-life adversity (same components). We included covariates known to influence survival and longevity (e.g., mother's age at first reproduction, lifetime density and food availability (lifetime spruce cones), study area, and whether she experienced a mast. Second, to test whether maternal adversity predicted daughter's lifetime fitness (LRS), we built a Poisson GLM with a log link including the same early-life adversity predictors for both generations. We also controlled for reproductive tenure and environmental quality across the period of reproduction. For both models, we assessed multicollinearity using variance inflation factors (VIFs).

### Supplementary Figures

**Fig. S1. Correlations among cohort and intrinsic sources of early-life adversity.** Heatmap shows pairwise Pearson correlation coefficients among the six birth-year sources of adversity: natal squirrel density, lynx–hare cycle phase at birth, birth date (standardized within study area and year), litter size (standardized within study area and year), presence/absence of a mast event, and postnatal growth rate (standardized within study area and year). Values within cells reflect Pearson's *r*, with darker shades indicating stronger positive (black) or negative (grey) correlations. Only the lower triangle is shown. Categorical variables were transformed prior to analysis: lynx–hare phase was converted to an ordered numeric factor and natal mast was coded as 0/1 (no/yes). Correlations were calculated using complete pairwise observations.

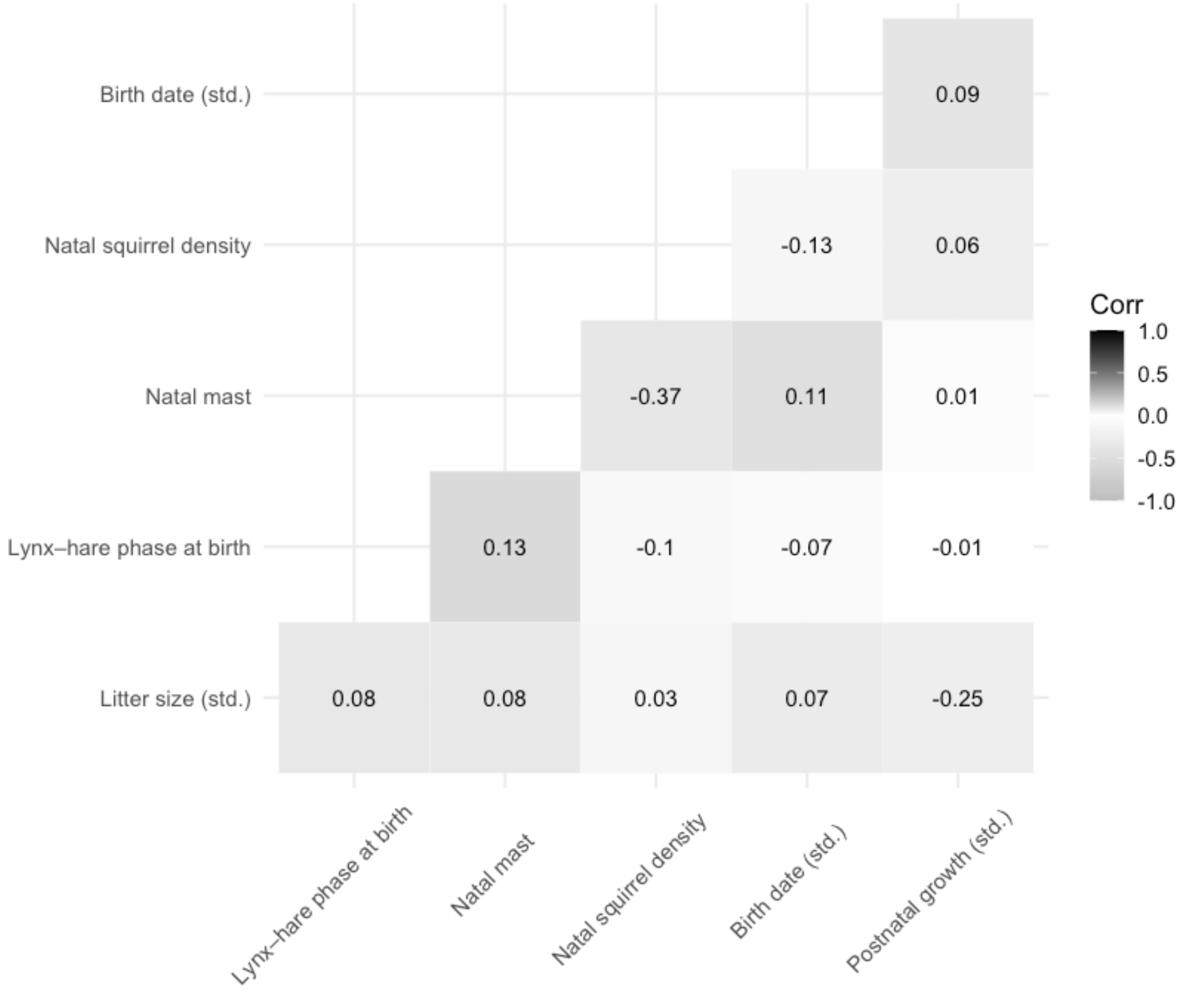

**Fig. S2. Variation in adult lifespan and reproductive tenure across the dataset (N=405 females).**

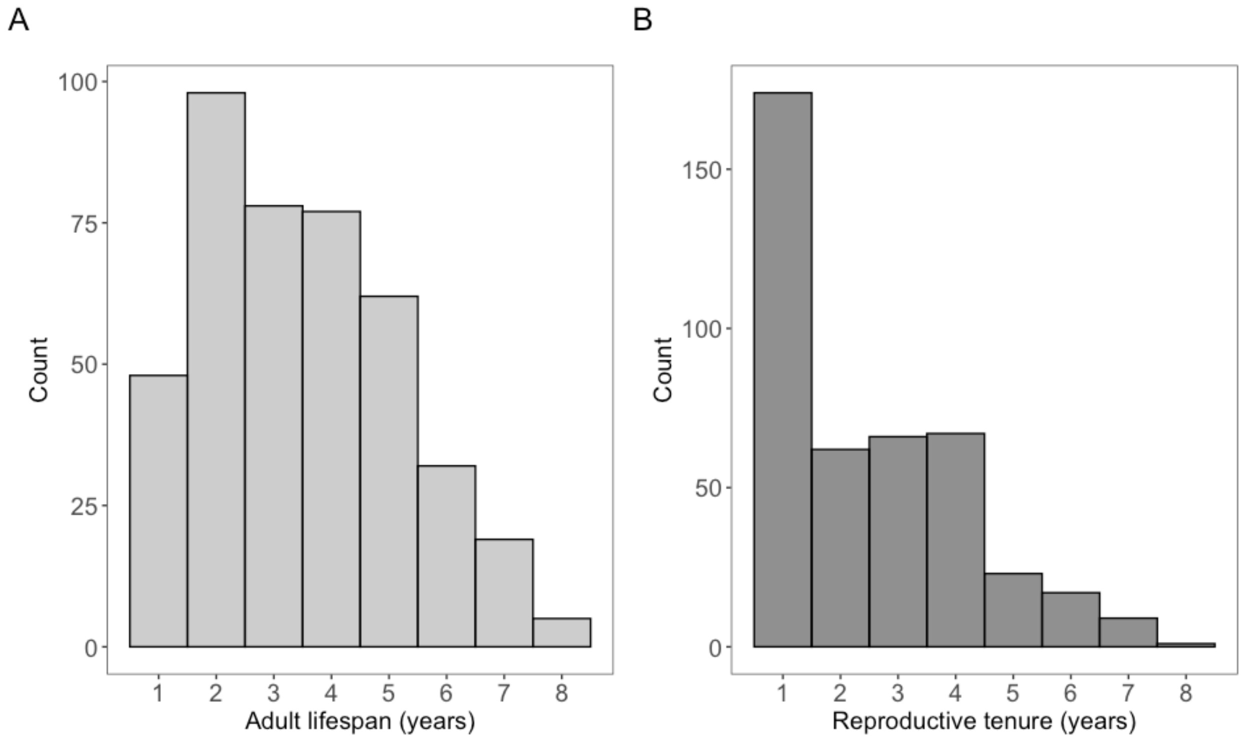

### Supplementary tables

**Table S1. Preliminary model quantifying sources of early-life adversity in our study population.** We included variables expected to influence variation in the probability of juvenile overwinter survival (>200 days of age, the main survival bottleneck in this population, (*37*)) based on prior support in this and other populations (e.g., (*36*)). Results from a generalized linear mixed-effects model (family=binomial) that included mother ID, year, and litter ID as random effects. We then extracted coefficients of significant predictors in this model to construct total, intrinsic, and cohort adversity indices ELA indices.

| **Juvenile overwinter survival (N=3,689 squirrels)** | | | | |
|---|---|---|---|---|
| | **Estimate** | **Std. error** | **z** | **P** |
| study area (SU) | 0.048 | 0.110 | 0.439 | 0.661 |
| **sex (male)** | **-0.767** | **0.097** | **-7.895** | **0.000** |
| **litter size** | **-0.156** | **0.066** | **-2.370** | **0.018** |
| **birth date** | **-0.263** | **0.058** | **-4.502** | **0.000** |
| **postnatal growth** | **0.282** | **0.056** | **5.046** | **0.000** |
| **mast (yes)** | **1.290** | **0.270** | **4.785** | **0.000** |
| **squirrel density** | **-0.499** | **0.118** | **-4.233** | **0.000** |
| hare-lynx cycle (peak) | 0.223 | 0.318 | 0.703 | 0.482 |
| **hare-lynx-cycle (year following peak)** | **-0.760** | **0.333** | **-2.284** | **0.022** |
| hare-lynx-cycle (2 years following peak) | -0.319 | 0.331 | -0.966 | 0.334 |
| mustelid abundance | 0.063 | 0.107 | 0.587 | 0.557 |
| mean overwinter temperature | -0.028 | 0.106 | -0.260 | 0.794 |
| **litter size x mast (yes)** | **0.355** | **0.117** | **3.028** | **0.002** |
| **birth date x squirrel density** | **-0.157** | **0.075** | **-2.104** | **0.035** |
| postnatal growth x squirrel density | -0.094 | 0.065 | -1.457 | 0.145 |

**Table S2. Longevity, but not early-life adversity, predicts variation in total lifetime fitness.** Neither intrinsic nor cohort adversity explained variation in lifetime reproductive output or lifetime reproductive success (LRS). In contrast, adult lifespan was a strong positive predictor of both metrics, indicating that overall longevity—rather than early-life adversity—drives differences in total reproductive performance. Table S4A shows results for lifetime reproductive output; Table S4B shows results for LRS. Results from generalized linear models (family=Poisson); Model B included an offset for total lifetime reproductive output.

| | **A) Lifetime reproductive output (N=405)** | | | | **B) Lifetime reproductive success (N=405)** | | | |
|---|---|---|---|---|---|---|---|---|
| | **Estimate** | **Std. error** | **z** | **P** | **Estimate** | **Std. error** | **z** | **P** |
| intrinsic adversity | -0.029 | 0.017 | -1.704 | 0.089 | -0.022 | 0.035 | -0.618 | 0.537 |
| cohort adversity | -0.009 | 0.019 | -0.498 | 0.619 | -0.003 | 0.039 | -0.068 | 0.946 |
| **adult lifespan** | **0.513** | **0.036** | **14.055** | **0.000** | **0.146** | **0.069** | **2.109** | **0.035** |
| **lifetime squirrel density** | **-0.185** | **0.031** | **-5.928** | **0.000** | **-0.195** | **0.061** | **-3.185** | **0.001** |
| lifetime spruce cones | 0.008 | 0.028 | 0.281 | 0.778 | 0.106 | 0.056 | 1.911 | 0.056 |
| **experienced mast (yes)** | **0.455** | **0.047** | **9.777** | **0.000** | **0.755** | **0.114** | **6.592** | **0.000** |
| **study area (SU)** | **-0.062** | **0.040** | **-1.562** | **0.118** | **0.207** | **0.080** | **2.569** | **0.010** |

**Table S3. Developmental adversity reduces lifespan.** Among females that survived beyond 200 days, higher rates of developmental adversity predicted shorter adult lifespans. This reduction was attributable to cohort adversity—reflecting harsh or competitive natal environments—rather than intrinsic adversity. These results are consistent with prior evidence in this and other species showing that early-life adversity shortens adult lifespan. Table S2A shows effects of total developmental adversity; Table S2B partitions total adversity into intrinsic and cohort components. Results from generalized linear models (family=gaussian).

| **A) Adult lifespan, years (N=405 females)** | | | | |
|---|---|---|---|---|
| | **Estimate** | **Std. error** | **z** | **P** |
| **total developmental adversity** | **-0.044** | **0.014** | **-3.022** | **0.003** |
| **age at first reproduction** | **0.072** | **0.016** | **4.393** | **0.000** |
| **lifetime squirrel density** | **0.284** | **0.021** | **13.829** | **0.000** |
| **lifetime spruce cones** | **0.165** | **0.023** | **7.295** | **0.000** |
| **experienced mast (yes)** | **0.174** | **0.032** | **5.512** | **0.000** |
| **study area (SU)** | **0.114** | **0.029** | **3.912** | **0.000** |

| **B) Adult lifespan, years (N=405 females)** | | | | |
|---|---|---|---|---|
| | **Estimate** | **Std. error** | **z** | **P** |
| intrinsic adversity | 0.006 | 0.013 | 0.504 | 0.615 |
| **cohort adversity** | **-0.057** | **0.015** | **-3.796** | **0.000** |
| **age at first reproduction** | **0.076** | **0.016** | **4.651** | **0.000** |
| **lifetime squirrel density** | **0.289** | **0.021** | **14.083** | **0.000** |
| **lifetime spruce cones** | **0.154** | **0.023** | **6.714** | **0.000** |
| **experienced mast (yes)** | **0.194** | **0.033** | **5.953** | **0.000** |
| **study area (SU)** | **0.107** | **0.029** | **3.676** | **0.000** |

**Table S4. Cohort adversity increases mortality risk across the life course.** Females that experienced higher rates of cohort adversity during development faced significantly higher hazards of death as adults (HR = 1.15, 95% CI [1.03–1.26]), indicating reduced survival prospects throughout life. In contrast, intrinsic adversity had no detectable effect on mortality risk. Table shows results from a Cox proportional hazards model predicting time to death.

| **Time to death (N=405 females)** | | | | | |
|---|---|---|---|---|---|
| | **Estimate** | **Estimate(exp)** | **Std. error** | **z** | **P** |
| intrinsic adversity | -0.007 | 0.993 | 0.051 | -0.135 | 0.892 |
| **cohort adversity** | **0.138** | **1.148** | **0.056** | **2.449** | **0.014** |
| **lifetime squirrel density** | **-1.209** | **0.299** | **0.097** | **-12.427** | **0.000** |
| **lifetime spruce cones** | **-0.714** | **0.490** | **0.097** | **-7.337** | **0.000** |
| **experienced mast (yes)** | **-0.836** | **0.434** | **0.130** | **-6.447** | **0.000** |
| **study area (SU)** | **-0.664** | **0.515** | **0.124** | **-5.342** | **0.000** |

**Table S5. Developmental adversity does not predict the likelihood that a female accrues any fitness across her life course.** Neither cohort nor intrinsic adversity influenced the probability of achieving any lifetime reproductive success (LRS > 0). Roughly one-third of females produced no surviving offspring, but the likelihood of reproduction was instead driven by lifespan and environmental factors such as mast years and resource availability. Table shows results from a binomial generalized linear mixed-effects model predicting whether females accrued any LRS during their lifetime. Results from a generalized linear model (family=binomial).

| **Lifetime reproductive success, LRS > 0 (N=405)** | | | | |
|---|---|---|---|---|
| | **Estimate** | **Std. error** | **z** | **P** |
| intrinsic adversity | -0.106 | 0.152 | -0.701 | 0.483 |
| cohort adversity | -0.018 | 0.124 | -0.144 | 0.886 |
| **adult lifespan** | **1.343** | **0.344** | **3.899** | **0.000** |
| lifetime squirrel density | 0.101 | 0.056 | 1.786 | 0.074 |
| **lifetime spruce cones** | **-0.225** | **0.069** | **-3.270** | **0.001** |
| **experienced mast (yes)** | **1.542** | **0.312** | **4.951** | **0.000** |
| **study area (SU)** | **0.692** | **0.286** | **2.425** | **0.015** |

**Table S6. Preserving longevity does not preserve lifetime fitness for females that experienced high rates of intrinsic adversity.** Although females that experienced high intrinsic adversity did not suffer reduced lifetime reproductive success (LRS), this was not explained by their apparent ability to maintain total lifespan. Table shows results from a generalized linear model (family=Poisson) predicting lifetime reproductive success and included an offset for total reproductive output.

| **Lifetime reproductive success (LRS, N=405)** | | | | |
|---|---|---|---|---|
| | **Estimate** | **Std. error** | **z** | **P** |
| intrinsic adversity | -0.011 | 0.040 | -0.267 | 0.790 |
| **adult lifespan** | **0.143** | **0.069** | **2.058** | **0.040** |
| cohort adversity | -0.001 | 0.039 | -0.036 | 0.972 |
| **lifetime squirrel density** | **-0.196** | **0.061** | **-3.191** | **0.001** |
| **lifetime spruce cones** | **0.109** | **0.056** | **1.950** | **0.051** |
| **experienced mast (yes)** | **0.757** | **0.115** | **6.605** | **0.000** |
| **study area (SU)** | **0.207** | **0.080** | **2.567** | **0.010** |
| intrinsic adversity x adult lifespan | -0.018 | 0.032 | -0.569 | 0.570 |

**Table S7. Reproductive tenure is a stronger predictor of lifetime fitness than adult lifespan.** The number of years a female reproduced (reproductive tenure) explained more variation in both lifetime reproductive output and lifetime reproductive success than adult lifespan did. Models including reproductive tenure were a better fit (ΔAICc ≈ –91 for output; –17 for success), indicating that the duration of active reproduction—not simply survival—best captures the component of lifespan that contributes to fitness. Results from generalized linear models (family=Poisson). Tables S7A–B compare models predicting lifetime reproductive output, and Tables S7C–D compare models predicting lifetime reproductive success.

**A) Lifetime reproductive output**
(N=405, AICc=2039.1)

| | Estimate | SE | z | P |
|---|---|---|---|---|
| **reproductive tenure** | **0.387** | **0.023** | **16.981** | **0.000** |
| lifetime spruce cones | 0.022 | 0.027 | 0.846 | 0.398 |
| lifetime squirrel density | -0.037 | 0.025 | -1.478 | 0.139 |
| **experienced mast (yes)** | **0.485** | **0.043** | **11.275** | **0.000** |
| study area (SU) | -0.010 | 0.038 | -0.248 | 0.804 |

**B) Lifetime reproductive output**
(N=405, AICc=2130.4)

| | Estimate | SE | z | P |
|---|---|---|---|---|
| **adult lifespan** | **0.519** | **0.036** | **14.323** | **0.000** |
| lifetime spruce cones | 0.009 | 0.028 | 0.336 | 0.737 |
| **lifetime squirrel density** | **-0.188** | **0.030** | **-6.211** | **0.000** |
| **experienced mast (yes)** | **0.451** | **0.043** | **10.491** | **0.000** |
| study area (SU) | -0.063 | 0.039 | -1.610 | 0.107 |

**C) Lifetime reproductive success**
(N=405, AICc=1479.9)

| | | | | |
|---|---|---|---|---|
| **reproductive tenure** | **0.447** | **0.045** | **9.912** | **0.000** |
| **lifetime spruce cones** | **0.140** | **0.052** | **2.692** | **0.007** |
| **lifetime squirrel density** | **-0.183** | **0.050** | **-3.674** | **0.000** |
| **experienced mast (yes)** | **1.290** | **0.109** | **11.889** | **0.000** |
| **study area (SU)** | **0.240** | **0.078** | **3.062** | **0.002** |

**D) Lifetime reproductive success**
(N=405, AICc=1496.7)

| | | | | |
|---|---|---|---|---|
| **adult lifespan** | **0.638** | **0.069** | **9.237** | **0.000** |
| **lifetime spruce cones** | **0.115** | **0.054** | **2.135** | **0.033** |
| **lifetime squirrel density** | **-0.366** | **0.060** | **-6.154** | **0.000** |
| **experienced mast (yes)** | **1.224** | **0.109** | **11.280** | **0.000** |
| **study area (SU)** | **0.159** | **0.081** | **1.978** | **0.048** |

**Table S8. Developmental adversity shortens reproductive tenure.** Females that experienced higher rates of developmental adversity had shorter reproductive tenures after accounting for differences in total adult lifespan. Both intrinsic and cohort adversity independently reduced the duration of reproduction over the life course, suggesting that early-life challenges constrain the ability to sustain lifetime reproductive effort. Table S8A shows effects of total developmental adversity; Table S8B separates intrinsic and cohort components. Both models included an offset for total adult lifespan.

**A) Reproductive tenure (offset by total lifespan, N=405)**

| | Estimate | Std. error | z | P |
|---|---|---|---|---|
| **total developmental adversity** | **-0.065** | **0.019** | **-3.365** | **0.001** |
| squirrel density across reproduction | -0.100 | 0.100 | -0.990 | 0.323 |
| **spruce cones across reproduction** | **0.210** | **0.099** | **2.118** | **0.035** |
| experienced mast (yes) | 0.035 | 0.039 | 0.905 | 0.366 |
| study area (SU) | -0.021 | 0.042 | -0.501 | 0.617 |

**B) Reproductive tenure (offset by total lifespan, N=405)**

| | Estimate | Std. error | z | P |
|---|---|---|---|---|
| **intrinsic adversity** | **-0.052** | **0.019** | **-2.781** | **0.006** |
| **cohort adversity** | **-0.047** | **0.020** | **-2.385** | **0.018** |
| squirrel density across reproduction | -0.126 | 0.101 | -1.241 | 0.215 |
| **spruce cones across reproduction** | **0.232** | **0.100** | **2.333** | **0.020** |
| experienced mast (yes) | 0.026 | 0.039 | 0.669 | 0.504 |
| study area (SU) | -0.015 | 0.042 | -0.360 | 0.719 |

**Table S9. An earlier age at first reproduction and a later age at last reproduction enhance lifetime fitness.** Females that began reproducing at younger ages achieved higher lifetime reproductive output and greater lifetime reproductive success (LRS), even after accounting for lifespan and environmental variation. These results indicate strong selection favoring earlier reproductive maturity. Table S9A shows effects on lifetime reproductive output; Table S9B shows effects on LRS. Results from generalized linear models (family=Poisson); Model B included an offset for total reproductive output.

**A) Dependent variable: Lifetime reproductive output (N=405)**

| | **Estimate** | **Std. error** | **z** | **P** |
|---|---|---|---|---|
| age at last reproduction | 0.331 | 0.038 | 8.623 | 0.000 |
| age at first reproduction | -0.155 | 0.020 | -7.674 | 0.000 |
| adult lifespan | 0.120 | 0.039 | 3.068 | 0.002 |
| spruce cones across reproduction | -0.082 | 0.104 | -0.793 | 0.428 |
| squirrel density across reproduction | 0.036 | 0.104 | 0.341 | 0.733 |
| experienced mast (yes) | 0.414 | 0.047 | 8.893 | 0.000 |
| study area (SU) | -0.045 | 0.042 | -1.060 | 0.289 |

**B) Dependent variable: Lifetime reproductive success (N=405)**

| | **Estimate** | **Std. error** | **z** | **P** |
|---|---|---|---|---|
| age at last reproduction | 0.238 | 0.079 | 3.000 | 0.003 |
| age at first reproduction | -0.099 | 0.043 | -2.282 | 0.022 |
| adult lifespan | 0.208 | 0.080 | 2.586 | 0.010 |
| spruce cones across reproduction | -0.136 | 0.195 | -0.696 | 0.486 |
| squirrel density across reproduction | 0.421 | 0.203 | 2.070 | 0.038 |
| experienced mast (yes) | 1.285 | 0.115 | 11.137 | 0.000 |
| study area (SU) | 0.079 | 0.089 | 0.892 | 0.373 |

**Table S10. Cohort adversity delays reproductive maturity.** Females that experienced high rates of developmental adversity reproduced later in life, contrary to expectations of accelerated reproductive maturity. This delay was driven entirely by cohort adversity, with age at first reproduction increasing by roughly two months for each standard deviation rise in cohort adversity. Table S10A shows effects of total developmental adversity; Table S10B separates intrinsic and cohort components. Results from generalized linear models (family=Poisson). Both models include year of first reproduction as a random effect.

**A) Age at first reproduction, AFR (N=405)**

| | Estimate | Std. error | z | P |
|---|---|---|---|---|
| **total developmental adversity** | **0.193** | **0.057** | **3.369** | **0.001** |
| **adult lifespan** | **0.165** | **0.052** | **3.197** | **0.001** |
| **mast at AFR** | **-0.341** | **0.172** | **-1.984** | **0.047** |
| **squirrel density at AFR** | **0.222** | **0.109** | **2.033** | **0.042** |
| spruce cones at AFR | -0.034 | 0.046 | -0.743 | 0.458 |
| study area (SU) | -0.052 | 0.111 | -0.469 | 0.639 |

**B) Age at first reproduction, AFR (N=405)**

| | Estimate | Std. error | z | P |
|---|---|---|---|---|
| intrinsic adversity | 0.004 | 0.047 | 0.080 | 0.936 |
| **cohort adversity** | **0.247** | **0.063** | **3.932** | **0.000** |
| **adult lifespan** | **0.171** | **0.052** | **3.305** | **0.001** |
| **mast at AFR** | **-0.357** | **0.173** | **-2.060** | **0.039** |
| **squirrel density at AFR** | **0.205** | **0.111** | **1.853** | **0.064** |
| spruce cones at AFR | -0.016 | 0.048 | -0.329 | 0.742 |
| study area (SU) | -0.043 | 0.112 | -0.384 | 0.701 |

**Table S11. Intrinsic adversity accelerates reproductive stop.** Females that experienced high rates of intrinsic adversity ceased reproduction at younger ages than their less-stressed counterparts, even after accounting for variation in total lifespan and environmental conditions. This effect reflects earlier reproductive cessation rather than earlier maturation, suggesting that physiological adversity accelerates reproductive senescence. Table shows results from a linear mixed-effects model predicting age at last reproduction (N = 405), with year of last reproduction included as a random effect.

| **Age at last reproduction, ALR (N=405)** | | | | | |
|---|---|---|---|---|---|
| | **Estimate** | **Std. error** | **df** | **t** | **P** |
| **intrinsic adversity** | **-0.111** | **0.036** | **375.419** | **-3.104** | **0.002** |
| cohort adversity | 0.030 | 0.043 | 393.789 | 1.589 | 0.515 |
| **adult lifespan** | **1.474** | **0.046** | **387.291** | **30.377** | **0.000** |
| **age at first reproduction** | **0.140** | **0.040** | **392.481** | **4.025** | **0.000** |
| **experienced mast (yes)** | **0.299** | **0.093** | **378.239** | **1.677** | **0.006** |
| lifetime squirrel density | -0.138 | 0.013 | 394.393 | -0.575 | 0.157 |
| lifetime spruce cones | 0.047 | 0.011 | 394.299 | 1.138 | 0.646 |
| **study area (SU)** | **-0.140** | **0.097** | **304.991** | **-1.448** | **0.148** |

**Table S12. Cohort adversity predicts frontloading of reproduction into the first reproductive attempt.** Females that experienced high rates of cohort adversity produced more offspring and successfully recruited more of those offspring in their first breeding season than females with lower rates of cohort adversity, even after accounting for environmental conditions at reproduction. This pattern indicates that cohort adversity favors early allocation of reproductive effort—frontloading reproduction into the first season—as a compensatory strategy to offset anticipated constraints on future reproductive opportunities. Table S12A shows effects on first-season reproductive output (number of offspring produced); Table S12B shows effects on first-season reproductive success (number of offspring surviving >200 days). Both models include a random effect of first-season year.

**A) Reproductive output in first breeding season (N=405 females)**

| | **Estimate** | **Std. error** | **z** | **P** |
|---|---|---|---|---|
| **cohort adversity** | **0.130** | **0.034** | **3.864** | **0.000** |
| intrinsic adversity | -0.009 | 0.026 | -0.337 | 0.736 |
| **age** | **0.247** | **0.028** | **8.745** | **0.000** |
| study area (SU) | 0.045 | 0.059 | 0.772 | 0.440 |
| **first-season mast** | **0.437** | **0.118** | **3.700** | **0.000** |
| **first-season squirrel density** | **-0.193** | **0.075** | **-2.590** | **0.010** |
| first-season spruce cones | 0.010 | 0.032 | 0.297 | 0.766 |

**B) Reproductive success in first breeding season (N=405 females, offset by reproductive output)**

| | | | | |
|---|---|---|---|---|
| **cohort adversity** | **0.285** | **0.087** | **3.280** | **0.001** |
| intrinsic adversity | -0.026 | 0.059 | -0.434 | 0.664 |
| **age at first reproduction** | **0.259** | **0.071** | **3.642** | **0.000** |
| **study area (SU)** | **0.430** | **0.229** | **1.879** | **0.060** |
| **first-season output** | **0.204** | **0.045** | **4.508** | **0.000** |
| **first-season mast** | **0.634** | **0.256** | **2.478** | **0.013** |
| first-season squirrel density | -0.205 | 0.131 | -1.565 | 0.118 |
| first-season spruce cones | 0.176 | 0.128 | 1.371 | 0.170 |

**Table S13. Cohort adversity increases the likelihood of concentrating lifetime fitness at debut.** Females that experienced high rates of cohort adversity were more likely to accrue their entire lifetime reproductive success (LRS) in their first breeding season, even after controlling for lifespan. Among females that continued breeding across multiple years, those from high-adversity cohorts also devoted a greater proportion of their total LRS to their first season. Results from generalized linear mixed-effects models (A) shows the binary outcome of achieving all LRS in the first season; B) shows the proportion of total LRS accrued at debut). Both models include a random effect of first-season year.

**A) Probability of accruing all LRS in first season (N=242 females, family=binomial)**

| | Estimate | Std. error | z | P |
|---|---|---|---|---|
| **cohort adversity** | **0.528** | **0.236** | **2.237** | **0.025** |
| intrinsic adversity | 0.064 | 0.173 | 0.370 | 0.711 |
| **first-season mast** | **1.117** | **0.403** | **2.773** | **0.006** |
| **first-season squirrel density** | **0.867** | **0.424** | **2.044** | **0.041** |
| first-season spruce cones | 0.264 | 0.145 | 1.825 | 0.068 |
| **adult lifespan** | **-1.564** | **0.276** | **-5.671** | **0.000** |
| experienced mast (yes) | -0.602 | 0.484 | -1.245 | 0.213 |
| spruce cones across reproduction | 0.060 | 0.071 | 0.837 | 0.403 |
| squirrel density across reproduction | -0.104 | 0.081 | -1.290 | 0.197 |
| study area (SU) | 0.641 | 0.416 | 1.540 | 0.123 |

**B) Proportion of LRS in first season (N=242 females, family=binomial)**

| | Estimate | Std. error | t | P |
|---|---|---|---|---|
| **cohort adversity** | **0.710** | **0.154** | **4.625** | **0.000** |
| intrinsic adversity | -0.038 | 0.096 | -0.396 | 0.692 |
| **first-season mast** | **1.224** | **0.482** | **2.538** | **0.011** |
| first-season squirrel density | -0.067 | 0.400 | -0.168 | 0.866 |
| first-season spruce cones | 0.110 | 0.135 | 0.813 | 0.416 |

| | | | | |
|---|---|---|---|---|
| adult lifespan | -0.098 | 0.150 | -0.652 | 0.514 |
| experienced mast (yes) | 0.555 | 0.730 | 0.760 | 0.447 |
| spruce cones across reproduction | -0.763 | 0.731 | -1.044 | 0.296 |
| squirrel density across reproduction | -0.388 | 0.323 | -1.201 | 0.230 |
| **lifetime reproductive output** | **-0.804** | **0.137** | **-5.872** | **0.000** |
| study area (SU) | 0.455 | 0.246 | 1.846 | 0.065 |

**Table S14. Frontloading reproduction offsets the fitness costs associated with cohort adversity, revealing fitness costs for those that do not frontload.** Females that experienced high rates of cohort adversity accrued greater lifetime reproductive success (LRS) when they frontloaded reproduction into their first breeding season. By contrast, cohort adversity predicted lower LRS among females that did not frontload (independent effect of cohort adversity, β=-0.12, P=0.004), revealing that early reproductive investment can mitigate the fitness costs of harsh developmental conditions. Table shows results from a generalized linear model (family=Poisson) predicting variation in lifetime reproductive success (N = 405 females).

| **Lifetime reproductive success, LRS (N=405 females, offset = reproductive lifespan)** | | | | |
|---|---|---|---|---|
| | **Estimate** | **Std. error** | **t** | **P** |
| **cohort adversity** | **-0.126** | **0.042** | **-3.016** | **0.003** |
| **first-season ARS** | **0.362** | **0.034** | **10.767** | **0.000** |
| intrinsic adversity | 0.006 | 0.039 | 0.151 | 0.880 |
| **first-season mast** | **-0.512** | **0.092** | **-5.565** | **0.000** |
| first-season spruce cones | -0.048 | 0.028 | -1.724 | 0.085 |
| **first-season squirrel density** | **-0.517** | **0.096** | **-5.380** | **0.000** |
| age at first reproduction | -0.092 | 0.051 | -1.799 | 0.072 |
| **squirrel density across reproduction** | **1.116** | **0.244** | **4.569** | **0.000** |
| **spruce cones across reproduction** | **-1.017** | **0.237** | **-4.282** | **0.000** |
| **experienced mast (yes)** | **0.884** | **0.106** | **8.375** | **0.000** |
| **study area (SU)** | **-0.247** | **0.088** | **-2.790** | **0.005** |
| **cohort adversity x first-season ARS** | **0.088** | **0.042** | **2.101** | **0.036** |
| intrinsic adversity x first-season ARS | -0.025 | 0.019 | -1.272 | 0.203 |

**Table S15. Developmental adversity shapes individual generation times.** Cohort adversity was associated with accelerated reproductive pace, whereas intrinsic adversity predicted later reproductive timing after controlling for age at first reproduction, reproductive tenure, lifetime spruce cone production, lifetime squirrel density, experienced mast, and study area. Table shows results from a generalized linear model (Gamma family, log link) testing effects of early-life adversity on *individual generation time* (offspring-weighted mean age at reproduction; *N* = 405 females). Negative coefficients indicate shorter generation times (faster reproduction); positive coefficients indicate slower generation times. We calculated generation time as the offspring-weighted mean age at reproduction [see Materials and Methods, (*50*)].

| **Individual generation time (N=405 females)** | | | | |
|---|---|---|---|---|
| | **Estimate** | **Std. error** | **t** | **P** |
| **cohort adversity** | **-0.080** | **0.011** | **-7.331** | **0.000** |
| **intrinsic adversity** | **0.026** | **0.010** | **2.649** | **0.008** |
| study area (SU) | -0.011 | 0.021 | -0.537 | 0.591 |
| **reproductive tenure** | **0.323** | **0.020** | **16.390** | **0.000** |
| **age at first reproduction** | **0.346** | **0.013** | **27.149** | **0.000** |
| lifetime spruce cones | -0.025 | 0.019 | -1.318 | 0.188 |
| **lifetime squirrel density** | **0.046** | **0.020** | **2.245** | **0.025** |
| experienced mast (yes) | 0.021 | 0.029 | 0.739 | 0.461 |

**Table S16. Cohort adversity predicts a greater likelihood of encountering a mast, and a shorter time to mast.** Results of (**A)** a binomial generalized linear model testing whether early-life adversity predicts the probability that a female ever experiences a mast year, and **(B)** a Cox proportional hazards model testing effects of adversity on the timing of first mast encounter (N = 405 females). Positive coefficients indicate higher likelihood or faster occurrence of mast exposure. Females that experienced cohort adversity were significantly more likely to experience a mast and did so sooner, whereas intrinsic adversity showed no detectable effect. Adult lifespan and study area were included as covariates in both models.

**A) Probability of encountering a mast (N=405 females)**

| | Estimate | Std. error | z | P |
|---|---|---|---|---|
| **cohort adversity** | **1.440** | **0.200** | **7.217** | **0.000** |
| intrinsic adversity | -0.397 | 0.323 | -1.229 | 0.219 |
| **adult lifespan** | **0.959** | **0.103** | **9.286** | **0.000** |
| study area (SU) | -0.183 | 0.264 | -0.694 | 0.487 |

**B) Time to mast (N=405 females)**

| | Estimate | Estimate(exp) | Std. error | z | P |
|---|---|---|---|---|---|
| **cohort adversity** | **0.631** | **1.879** | **0.070** | **9.064** | **0.000** |
| intrinsic adversity | 0.017 | 1.017 | 0.066 | 0.264 | 0.792 |

**Table S17. Cohort adversity does not increase the probability of the first reproductive attempt coinciding with a mast year.** Neither cohort nor intrinsic adversity significantly influenced the likelihood of first reproducing during a mast, suggesting that females did not delay reproductive maturity until high-payoff conditions were detected. Table shows results of a binomial generalized linear model testing whether early-life adversity predicts the probability that a female's first reproductive attempt occurred in a mast year (N = 405 females). Study area was included as a covariate.

| **Probability of first reproductive attempt during a mast (N=405 females)** | | | | |
|---|---|---|---|---|
| | **Estimate** | **Std. error** | **z** | **P** |
| intrinsic adversity | -0.399 | 0.264 | -1.508 | 0.132 |
| cohort adversity | -0.169 | 0.133 | -1.274 | 0.203 |
| study area (SU) | -0.188 | 0.210 | -0.894 | 0.371 |

**Table S18. Frontloading of reproductive output persists under poor conditions.** Table shows results from generalized linear models testing whether early-life adversity predicts first-season reproductive output **(A)** and reproductive success **(B)** among females whose first reproductive attempt occurred only in a non-mast year (N=230). Cohort adversity predicts greater reproductive output at debut even when the first reproductive attempt was not in a mast year. All models included age at first reproduction, study area, and local environmental covariates (spruce cones and squirrel density at first reproduction).

| **A) Reproductive output (N=230 females)** | | | | |
|---|---|---|---|---|
| | **Estimate** | **Std. error** | **z** | **P** |
| **cohort adversity** | **0.103** | **0.049** | **2.116** | **0.034** |
| intrinsic adversity | -0.009 | 0.039 | -0.227 | 0.820 |
| **age at first reproduction (y)** | **0.212** | **0.039** | **5.459** | **0.000** |
| study area (SU) | 0.150 | 0.093 | 1.605 | 0.109 |
| density at first reproduction | -0.119 | 0.074 | -1.598 | 0.110 |
| spruce cones at first reproduction | 0.037 | 0.029 | 1.287 | 0.198 |
| **B) Reproductive success (N=230 females, offset by reproductive output)** | | | | |
| | **Estimate** | **Std. error** | **z** | **P** |
| cohort adversity | -0.187 | 0.111 | -1.677 | 0.094 |
| intrinsic adversity | 0.063 | 0.093 | 0.683 | 0.495 |
| age at first reproduction (y) | -0.077 | 0.082 | -0.943 | 0.346 |
| study area (SU) | 0.334 | 0.195 | 1.714 | 0.087 |
| density at first reproduction | -0.115 | 0.110 | -1.043 | 0.297 |
| spruce cones at first reproduction | -0.073 | 0.119 | -0.617 | 0.537 |

**Table S19. Cohort adversity amplifies the costs of frontloading reproduction.** Under heightened extrinsic mortality risk during early life, preferential investment in the first reproductive attempt trades off more sharply with longevity. Females that experienced high rates of cohort adversity suffered stronger lifespan reductions when they frontloaded reproduction into their first breeding season. In contrast, females that delayed or distributed reproductive effort more evenly across years lived longer. Tables shows results from a generalized linear model (family=Gamma) predicting variation in adult lifespan as well as reproductive tenure in years (N = 405 females).

**A) Adult lifespan (N=405 females)**

| | Estimate | Std. error | t | P |
|---|---|---|---|---|
| **cohort adversity** | **-0.065** | **0.015** | **-4.204** | **0.000** |
| first-season ARS | 0.027 | 0.017 | 1.565 | 0.118 |
| intrinsic adversity | 0.007 | 0.013 | 0.535 | 0.593 |
| **study area (SU)** | **0.101** | **0.029** | **3.447** | **0.001** |
| **age at first reproduction** | **0.061** | **0.014** | **4.305** | **0.000** |
| first-season output | 0.003 | 0.015 | 0.194 | 0.847 |
| **lifetime squirrel density** | **0.294** | **0.021** | **14.214** | **0.000** |
| **lifetime spruce cones** | **0.150** | **0.023** | **6.446** | **0.000** |
| **experienced mast (yes)** | **0.189** | **0.034** | **5.568** | **0.000** |
| **cohort adversity x first-season ARS** | **-0.039** | **0.019** | **-2.033** | **0.043** |
| intrinsic adversity x first-season ARS | -0.007 | 0.012 | -0.597 | 0.551 |

**B) Reproductive tenure (N=405 females)**

| | Estimate | Std. error | t | P |
|---|---|---|---|---|
| **cohort adversity** | **-0.128** | **0.031** | **-4.193** | **0.000** |
| first-season ARS | 0.006 | 0.035 | 0.184 | 0.854 |
| **intrinsic adversity** | **-0.048** | **0.027** | **-1.758** | **0.080** |
| study area (SU) | -0.105 | 0.062 | -1.702 | 0.090 |
| **age at first reproduction** | **-0.105** | **0.030** | **-3.551** | **0.000** |

| | | | | |
|---|---|---|---|---|
| first-season output | -0.073 | 0.031 | -2.313 | 0.021 |
| squirrel density across reproduction | 0.136 | 0.147 | 0.930 | 0.353 |
| spruce cones across reproduction | -0.041 | 0.145 | -0.283 | 0.777 |
| **experienced mast (yes)** | **0.718** | **0.060** | **11.992** | **0.000** |
| **cohort adversity x first-season ARS** | **-0.098** | **0.040** | **-2.482** | **0.013** |
| intrinsic adversity x first-season ARS | 0.009 | 0.025 | 0.353 | 0.724 |

**Table S20. A mast at first reproduction intensifies the lifespan penalty associated with cohort adversity and reproductive frontloading.** Table shows results from a generalized linear model testing whether reproductive context modifies the relationship between early-life adversity and adult lifespan (N = 405 females). The interaction between cohort adversity and mast at first reproduction was negative, indicating that females that experienced extrinsic mortality risk during development suffered steeper declines in lifespan when their first breeding attempt occurred during a mast year. Lifespan increased with age at first reproduction, lifetime resource abundance (spruce cones), and lower population density.

| **Adult lifespan (years, N=405 females)** | | | | |
|---|---|---|---|---|
| | **Estimate** | **Std. error** | **t** | **P** |
| cohort adversity | -0.029 | 0.019 | -1.501 | 0.134 |
| mast at first reproduction | -0.031 | 0.030 | -1.038 | 0.300 |
| reproductive success in first season | -0.015 | 0.029 | -0.515 | 0.607 |
| intrinsic adversity | -0.002 | 0.017 | -0.105 | 0.916 |
| **study area (SU)** | **0.096** | **0.029** | **3.269** | **0.001** |
| **age at first reproduction** | **0.081** | **0.019** | **4.360** | **0.000** |
| **lifetime squirrel density** | **0.300** | **0.020** | **14.738** | **0.000** |
| **lifetime spruce cones** | **0.149** | **0.024** | **6.333** | **0.000** |
| **experienced mast (yes)** | **0.201** | **0.035** | **5.826** | **0.000** |
| **cohort adversity x mast at first reproduction** | **-0.093** | **0.031** | **-2.965** | **0.003** |
| cohort adversity x success in first season | 0.021 | 0.029 | 0.738 | 0.461 |
| **mast at first season x success in first season** | **0.082** | **0.036** | **2.273** | **0.024** |
| mast at first season x intrinsic adversity | 0.028 | 0.027 | 1.003 | 0.316 |
| success in first season x intrinsic adversity | 0.013 | 0.025 | 0.512 | 0.609 |
| **cohort adversity x mast at first reproduction x success in first season** | **-0.100** | **0.039** | **-2.558** | **0.011** |
| intrinsic adversity x mast at first reproduction x success in first season | -0.032 | 0.029 | -1.111 | 0.267 |

**Table S21. Intrinsic adversity modulates a later-life rise in reproductive success characteristic of terminal investment.** Reproductive success (offspring survival rate per reproductive attempt; N = 1,075 reproductive attempts) increased as females neared death, consistent with terminal investment. This increase followed a curvilinear trajectory, peaking at intermediate proximity to death before declining in the final year of life. Females that experienced higher intrinsic adversity trended toward steeper increases. They exhibited more pronounced peaks and sharper post-peak declines, indicating that intrinsic adversity amplifies both the intensity and duration of terminal investment. Results from a generalized linear mixed-effects model (family = binomial) including squirrel ID and grid:year as random effects.

| **Reproductive success (rate of offspring survival in a given year, N=1075 reproductive attempts)** | | | | |
|---|---|---|---|---|
| | **Estimate** | **Std. error** | **z** | **P** |
| intrinsic adversity | -0.012 | 0.063 | -0.191 | 0.849 |
| **years from death** | **0.654** | **0.147** | **4.464** | **0.000** |
| **years from death (quad)** | **-0.479** | **0.143** | **-3.363** | **0.001** |
| cohort adversity | -0.047 | 0.070 | -0.669 | 0.504 |
| study area (SU) | 0.215 | 0.215 | 0.999 | 0.318 |
| **mast (yes)** | **0.983** | **0.230** | **4.266** | **0.000** |
| age | 0.047 | 0.071 | 0.672 | 0.502 |
| **squirrel density** | **-0.417** | **0.128** | **-3.261** | **0.001** |
| prior year food availability | 0.029 | 0.107 | 0.268 | 0.788 |
| **intrinsic adversity x years from death** | **0.271** | **0.140** | **1.936** | **0.053** |
| **intrinsic adversity x years from death (quad)** | **-0.300** | **0.132** | **-2.280** | **0.023** |
| cohort adversity x years from death | -0.131 | 0.148 | -0.883 | 0.377 |
| cohort adversity x years from death (quad) | 0.067 | 0.147 | 0.458 | 0.647 |

**Table S22. Increased output in the final breeding season compensates for decreased reproductive efficiency.** Despite reduced efficiency, females with high rates of intrinsic adversity increased reproductive output in their final breeding season maintained lifetime fitness. This indicates that terminal investment in offspring quantity rather than quality can preserve lifetime reproductive success (LRS) despite the developmental constraints associated with early-life intrinsic adversity. Results from a generalized linear model.

| **Lifetime reproductive output (N=405 females)** | | | | |
|---|---|---|---|---|
| | **Estimate** | **Std. error** | **z** | **P** |
| cohort adversity | -0.051 | 0.049 | -1.029 | 0.303 |
| last season reproductive output | -0.014 | 0.008 | -1.834 | 0.067 |
| **age at last reproduction** | **-0.210** | **0.033** | **-6.289** | **0.000** |
| intrinsic adversity | -0.092 | 0.052 | -1.773 | 0.076 |
| **lifetime reproductive output** | **0.055** | **0.007** | **7.457** | **0.000** |
| mast in last season | 0.025 | 0.080 | 0.306 | 0.759 |
| **spruce cones year before last season** | **0.060** | **0.029** | **2.100** | **0.036** |
| **density in last season** | **-0.197** | **0.069** | **-2.875** | **0.004** |
| study area (SU) | 0.083 | 0.086 | 0.963 | 0.336 |
| spruce cones across reproduction | -0.018 | 0.187 | -0.095 | 0.924 |
| squirrel density across reproduction | 0.163 | 0.200 | 0.816 | 0.415 |
| **experienced mast (yes)** | **1.122** | **0.124** | **9.045** | **0.000** |
| cohort adversity x last season output | 0.010 | 0.006 | 1.578 | 0.115 |
| **intrinsic adversity x last season output** | **0.019** | **0.008** | **2.293** | **0.022** |

**Table S23. Intrinsic adversity constrains the fitness benefits of reproductive success in the final breeding season.** Females that experienced high intrinsic adversity in the first year of life gained fewer recruits per offspring produced in their final breeding season. This reduced ability to convert reproductive output into realized success suggests physiological constraints that limit the fitness returns of late-life reproduction. Table shows results from a generalized linear model (family=Poisson).

| **Lifetime reproductive success, LRS (N = 405 females, offset = lifespan)** | | | | |
|---|---|---|---|---|
| | **Estimate** | **Std. error** | **z** | **P** |
| cohort adversity | 0.017 | 0.044 | 0.377 | 0.706 |
| **last season reproductive success** | **0.299** | **0.027** | **10.911** | **0.000** |
| **age at last reproduction** | **-0.193** | **0.034** | **-5.648** | **0.000** |
| intrinsic adversity | 0.022 | 0.042 | 0.541 | 0.589 |
| **lifetime reproductive output** | **0.062** | **0.007** | **8.304** | **0.000** |
| mast in last season | 0.168 | 0.083 | 2.028 | 0.043 |
| spruce cones year before last season | 0.062 | 0.029 | 2.116 | 0.034 |
| density in last season | -0.004 | 0.070 | -0.064 | 0.949 |
| **last season reproductive output** | **-0.271** | **0.043** | **-6.354** | **0.000** |
| **study area (SU)** | **0.195** | **0.086** | **2.272** | **0.023** |
| spruce cones across reproduction | 0.146 | 0.191 | 0.768 | 0.442 |
| squirrel density across reproduction | -0.079 | 0.201 | -0.392 | 0.695 |
| **experienced mast (yes)** | **0.917** | **0.125** | **7.366** | **0.000** |
| cohort adversity x last season reproductive success | -0.015 | 0.027 | -0.547 | 0.584 |
| **intrinsic adversity x last season reproductive success** | **-0.054** | **0.022** | **-2.461** | **0.014** |

**Table S24. Maternal intrinsic adversity reduced both the adult lifespan and LRS of daughters' lifespan and lifetime fitness.** Using data from N = 230 mother–offspring pairs, we tested whether intrinsic adversity experienced by mothers predicted daughters' **(A)** lifespan and **(B)** lifetime reproductive success (LRS), independent of the daughters' own developmental conditions. Lifespan was modeled with a Gamma GLM (log link), and LRS with a Poisson GLM (log link). Predictor variables included daughter and mother's intrinsic adversity, cohort adversity, age at first reproduction, density and spruce availability across the lifetime (A) and reproduction (B), whether they experienced a mast or not, and study area. There was no significant intergenerational effect of cohort adversity.

| **A) Daughter's adult lifespan (years, N=230)** | | | | |
|---|---|---|---|---|
| | **Estimate** | **Std. error** | **t** | **P** |
| intrinsic adversity | 0.007 | 0.018 | 0.393 | 0.694 |
| **mother's intrinsic adversity** | **-0.043** | **0.018** | **-2.421** | **0.016** |
| **cohort adversity** | **-0.075** | **0.021** | **-3.547** | **0.000** |
| mother's cohort adversity | 0.011 | 0.019 | 0.609 | 0.543 |
| **age at first reproduction** | **0.065** | **0.021** | **3.065** | **0.002** |
| **lifetime squirrel density** | **0.298** | **0.029** | **10.391** | **0.000** |
| **lifetime spruce cones** | **0.163** | **0.032** | **5.041** | **0.000** |
| **experienced mast (yes)** | **0.150** | **0.045** | **3.370** | **0.001** |
| **study area (SU)** | **0.156** | **0.041** | **3.797** | **0.000** |
| **B) Daughter's lifetime reproductive success (N=230)** | | | | |
| | **Estimate** | **Std. error** | **z** | **P** |
| intrinsic adversity | 0.056 | 0.048 | 1.155 | 0.248 |
| **mother's intrinsic adversity** | **-0.119** | **0.045** | **-2.621** | **0.009** |
| **reproductive tenure** | **0.483** | **0.050** | **9.651** | **0.000** |
| cohort adversity | -0.051 | 0.055 | -0.925 | 0.355 |
| mother's cohort adversity | 0.014 | 0.049 | 0.278 | 0.781 |
| age at first reproduction | 0.028 | 0.068 | 0.404 | 0.686 |
| squirrel density across reproduction | -0.102 | 0.263 | -0.389 | 0.697 |
| spruce cones across reproduction | 0.186 | 0.258 | 0.723 | 0.470 |
| **experienced mast (yes)** | **1.069** | **0.141** | **7.594** | **0.000** |
| study area (SU) | 0.057 | 0.105 | 0.543 | 0.587 |
